# Transmembrane Domain Dominance Drives Emergent Signaling and Allosteric Inversion in mGlu_1/5_ Heterodimers

**DOI:** 10.64898/2026.05.20.726619

**Authors:** Justin B. Steinfeld, Xia Lei, Madeline Laramee, Xin Lin, Alice L. Rodriguez, Paul K. Spearing, Wesley B. Asher, Colleen M. Niswender, Jonathan A. Javitch

## Abstract

Class C GPCRs function as obligate dimers in which only one G protein can engage the complex at a time, but how each protomer contributes to heterodimer coupling has remained unresolved. Using CODA-RET, a BRET-based assay reporting direct Gα_q_ recruitment to defined, full-length receptor pairs, we show that signaling at the mGlu₁/₅ heterodimer flows predominantly through the mGlu₁ protomer; domain-swapped chimeras localize this dominance to the transmembrane domain. The dominance generates emergent signaling: cis-acting mGlu₁ PAMs and NAMs undergo allosteric inversion when coupling is restricted to mGlu₅. By contrast, the mGlu₅-selective NAM MTEP is silent at the heterodimer, mirroring mGlu₅’s minimal role in driving Gα_q_. Because the mGlu₁ PAM tested acts only in cis, a trans-acting mGlu₁ PAM would theoretically be selective for mGlu₁/₁ homomers. These findings open a pharmacological design space in which protomer target and cis-versus-trans mode of action tune selectivity across mGlu₁/₁, mGlu₅/₅, and mGlu₁/₅ dimers.

## Introduction

Metabotropic glutamate receptors have long promised to be transformative drug targets for neuropsychiatric disorders, yet that promise remains largely unfulfilled.^1^ One obstacle is that these receptors function in the brain not as isolated entities, but as obligate dimers with complex signaling properties that remain poorly understood.^1^ Group I mGlu receptors, mGlu_1_ and mGlu_5_, are of particular therapeutic interest given their critical roles in neurodevelopment and synaptic plasticity.^2–7^ These receptors are expressed heterogeneously across brain regions and can form both mGlu_1/1_ and mGlu_5/5_ homodimers as well as mGlu_1/5_ heterodimers with potentially distinct signaling properties, raising the possibility that heterodimer-selective pharmacology could enable more precise circuit-level targeting.^8–11^

Importantly, mGlu receptor heterodimers can exhibit pharmacological properties distinct from their parent homodimers, with significant implications for drug development. For example, in the brain the effect of an mGlu_4_ selective positive allosteric modulators depend on whether it activates only mGlu_4/4_ homodimers or both homodimers and mGlu_2/4_ heterodimers.^12^ Notably, heterodimerization can change receptor function: mGlu_4/4_ homodimers support both cis and trans activation, whereas mGlu_2/4_ heterodimers are restricted to cis activation.^12^ Similarly, some mGlu_7_ negative allosteric modulators (NAMs) that block homodimers fail to block mGlu_7/8_ heterodimers.^13^ Thus, heterodimer pharmacology cannot be predicted from homodimer studies alone.

One previous study examining Group I heterodimers reported symmetric signal transduction, with both protomers contributing equally to G protein activation and capable of trans activation.^11^ However, this contradicts both previous and more recent work showing that both VFTs need to be bound for full activation of Group I mGlu receptors.^8,14–16^ Two methodological features of that study may account for this discrepancy: the constructs replaced native C-terminal domains with GABA_B_ tails, which is potentially problematic given that Group I mGlu receptor signaling is sensitive to C-terminal modifications^8,17–19^, and signaling was assessed using downstream calcium mobilization assays, where signal amplification could obscure differences in proximal receptor activation. Whether these findings—symmetric protomer contributions and functional trans activation—hold for Group I heterodimers with intact C-termini and more direct measures of receptor activation, and how allosteric modulators behave at these heterodimers, have remained unexplored.

Here we address these questions using CODA-RET (Complemented Donor Acceptor Resonance Energy Transfer), a bioluminescence-based approach that reports direct G protein recruitment to intact receptors retaining their full C-terminal domains, allowing us to distinguish signaling contributions from each protomer in a defined heterodimer.^20^ This approach reveals an unexpected asymmetry: within the mGlu_1/5_ heterodimer, Gα_q_ signaling occurs almost exclusively through the mGlu_1_ protomer. This dominance is mediated by the transmembrane domain and has striking consequences for allosteric pharmacology— mGlu_1_ positive allosteric modulators (PAMs) and NAMs that act in cis show inverted effects when Gα_q_ coupling is restricted to mGlu_5_ and the mGlu_5_ NAM, MTEP, is unable to modulate mGlu_1/5_. Additionally, we reproduce these CODA-RET findings using GABA_B_ tail assays similar to those used previously, but with a small linker between the truncated C-tail and the GABA_B_ tail, which likely relieves steric hindrance that impacted receptor function in prior studies. These findings suggest that heterodimers represent a distinct receptor population with unique mechanistic and pharmacological properties and raise the possibility that allosteric modulators with similar effects on homodimers may behave quite differently at heterodimers, depending on whether they act in cis or trans within the dimer.

## Results

### Design and validation of CODA-RET system

To measure Gα_q_ protein recruitment to Group I mGlu receptor dimers and distinguish the signaling contributions of each protomer, we adapted the CODA-RET system (Figure 1).^12,13,20^ In CODA-RET, complementary luciferase fragments are placed on specific protomers, so that Nanoluciferase (NLuc) reconstitutes only when defined receptor pairs—homodimers or heterodimers—dimerize, enabling BRET to mVenus-tagged mGα_sq_, an engineered mini-G protein, upon receptor activation.^21^

**Figure 1:**
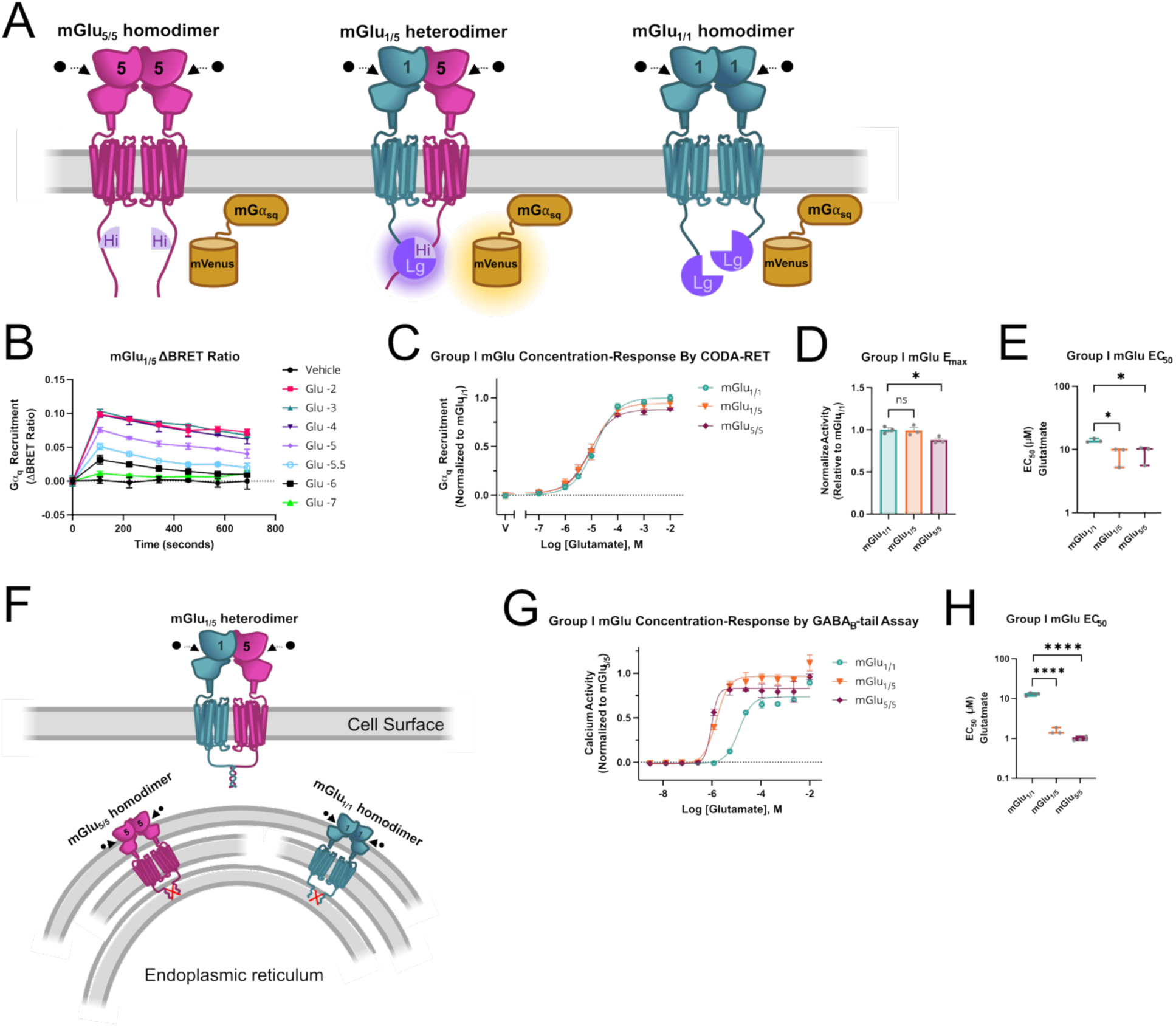
Heterodimer Activity Via CODA-RET and GABA_B_-tail Strategies. (A) Schematic of CODA-RET with mGlu_5_-HiBit homomer (left, magenta), mGlu_1_-LgBiT/mGlu_5_-HiBit heterodimer (middle, cyan/magenta), and mGlu_1_-LgBit homodimer (right, cyan). Glutamate is represented as black dots, binding into VFT’s and recruiting mVenus-mini-Gɑ_sq_ (gold), resulting in BRET only for the middle condition. (B) Example data from single experiment showing ΔBRET ratio for mGlu_1/5_ with vehicle and then decreasing amounts of glutamate (10^−2^ in pink to 10^−7^ in green) over time. Error bars represent ± SEM, each point is average of simultaneous triplicate measurement (3-wells) (C) Concentration-response curve for mGlu_1/1_ (cyan), mGlu_1/5_ (orange), mGlu_5/5_ (magenta) normalized to mGlu_1/1_ E_max_. Symbols represent the mean drug-induced BRET response and error bars represent ± SEM. (D) Corresponding E_max_ for Figure 1C. n.s. = 0.85, *p = 0.02. (E) Corresponding EC_50_ for Figure 1C. *p = 0.03 and *p = 0.04. (F) Schematic of GABA_B_-tail assay showing at the top that only mGlu_1_-GB1 and mGlu_5_-GB2 reaches the surface while at the bottom homodimers remain in the endoplasmic reticulum. (G) Concentration-response curve with glutamate and each homodimer (mGlu_1/1_ in cyan, mGlu_5/5_ in magenta) and heterodimer (mGlu_1/5_ in orange) normalized to mGlu_5/5_ E_max_. (H) Corresponding EC_50_ for Figure 1G. ****p < 0.0001 and ****p < 0.0001. Symbols represent the mean drug induced calcium-sensitive fluorescence and error bars represent ± SEM. The exact number of ‘n’ independent experiments and technical replicates are reported in Table S2.

Applying CODA-RET to Group I mGlu receptors required addressing a challenge posed by these receptors: their large C-terminal domains (>300 amino acids), which are critical for full receptor function.^8,17–19,22^ In previous CODA-RET studies with mGlu 2, 4, 7, and 8, split luciferases were placed at the receptor C-termini.^12,13,20^ This approach failed for Group I receptors—C-terminal probes produced little to no BRET signal (data not shown), likely because the distance from the membrane-proximal Gα_q_ to the distal C-terminus exceeds the optimal BRET range (∼55 Å).^23^ We therefore inserted an 11-amino-acid HiBiT sequence at an internal site within the C-terminal domains of these receptors, positioning it closer to the intracellular membrane. For mGlu_1_, we reasoned that the mGlu_1b_ splice site would tolerate probe insertion, since endogenous receptor variants naturally terminate at this position; for mGlu_5_, we identified a structurally equivalent site (Figures 1B and 1C; Table S1).^1^ We compared the Ca^2+^ mobilization activation of mGlu_1_ and mGlu_5_ wild type (WT) with their respective constructs with HiBiT insertions and LgBiT C-terminal additions and showed there was no discernible difference in their concentration-response curves, E_max_, or EC_50_ (Figure S1). Experiments in main figures used a single polarity, mGlu_1_-LgBiT and mGlu_5_-HiBiT; key experiments were repeated using the reverse polarity, mGlu_1_-HiBiT and mGlu_5_-LgBiT, which we designate mGlu_1/5_*, with results shown in Figure S5.

We performed two controls to validate the system using a GCaMP assay to measure downstream calcium activity. First, mutations at the G protein binding sites (mGlu_1_^F781D^; mGlu_5_^F767D^) or VFT orthosteric binding sites (mGlu_1_^T188A^; mGlu_5_^T174A^) within receptor protomers abolished agonist-induced calcium activity, confirming that signal depends on functional G protein coupling (Figure S1).^18,24–27^ Second, using CODA-RET, co-expression of excess wild-type (WT) untagged (“dark”) receptors (10x more DNA than the tagged receptors) with G protein binding-deficient luciferase-complemented receptors produced very little agonist-induced change in BRET, ruling out the possibility that G protein recruitment by nearby receptors could produce BRET through proximity to Nluc complemented from two signaling-dead protomers (Figure S2A). This confirmed that the agonist-induced BRET signal resulted from G protein coupling to the defined complemented dimer.

With this validated system, we characterized glutamate responses at mGlu_1/1_ and mGlu_5/5_ homodimers and the mGlu_1/5_ heterodimer. Maximal efficacy (E_max_) was equivalent for the mGlu_1/1_ homodimer and the mGlu_1/5_ heterodimer, but modestly and significantly lower at the mGlu_5/5_ homodimer (Figure 1D). Glutamate potency (EC_50_) was higher at the mGlu_5/5_ homodimer (8.8 ± 1.6 µM ) and mGlu_1/5_ heterodimer (8.4 ± 1.6 µM) than at the mGlu_1/1_ homodimer (13.9 ± 0.6 µM) (Figure 1E)—values consistent with previously published potencies for the homodimers.^28,29^ Luminescence levels were similar across conditions and did not correlate with ligand-induced BRET ratios, indicating that differences in receptor expression do not account for the observed signaling differences (Figure S2C).

To provide orthogonal validation, we also performed complementary experiments using GABA_B_-tail constructs with calcium mobilization assays. These constructs are similar to those used previously, with one critical difference that may account for the discrepancy in findings: a small AAA linker was added between the truncated C-tail of the mGlu receptor and the GABA_B_ tail, which may relieve steric hindrance that impacted native receptor function (Figures 1F and 1G). Again, the glutamate potency was higher at the mGlu_5/5_ homodimer (0.96 ± 0.05 µM) and the mGlu_1/5_ heterodimer (1.5 ± 0.2 µM) than at the mGlu_1/1_ homodimer (12.9 ± 0.6 µM), values in the same range as previously reported (Figures 1F, 1G, and 1H).^28,29^

### Asymmetric activation of the mGlu_1/5_ heterodimer

Previous work reported that mGlu_1/5_ heterodimers exhibit symmetric signal transduction.^11^ To test this with intact receptors, we systematically examined trans activation, cis activation, and protomer-specific signaling across Group I homodimers and heterodimers using both CODA-RET (Figures 2-4) and GABA_B_-tail calcium mobilization assays (Figures 4E and S3).

**Figure 2.**
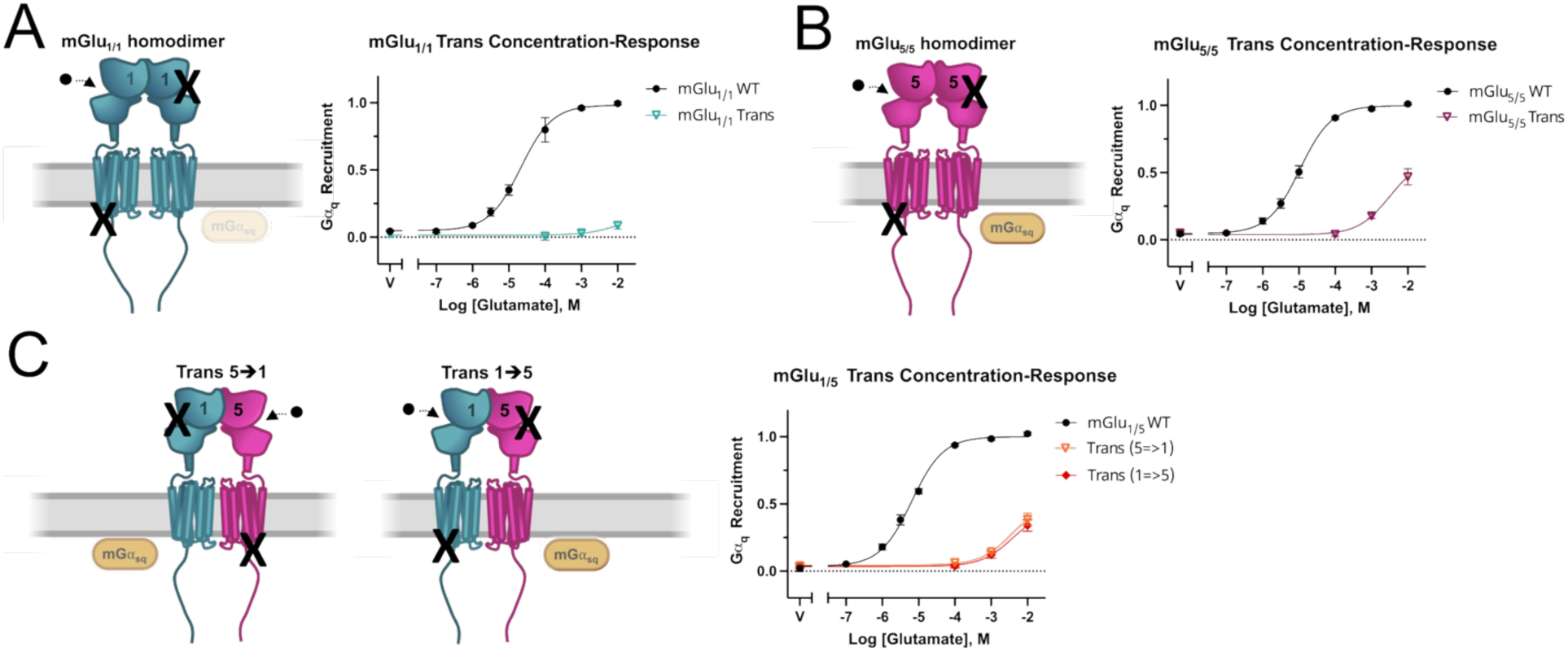
Group I mGlu Receptors Demonstrate Poor Trans Activation. (A) Left shows schematic of T188A and F781D mutation on each mGlu_1_ protomer (cyan), restricting to transactivation. Right shows a concentration-response curve of mGlu_1/1_ WT (black) versus mGlu_1/1_ in trans (cyan) by CODA-RET. (B) Left shows schematic of T174A and F767D mutation on each mGlu_5_ protomer (magenta), restricting to transactivation. Right shows a concentration-response curve of mGlu_5/5_ WT (black) versus mGlu_5/5_ in trans (magenta) by CODA-RET. (C) Left shows schematic of T188A mutation on mGlu_1_ protomer (cyan) and F767D mutation on mGlu_5_ protomer (magenta), restricting to trans activation from mGlu_5_ to mGlu_1_. Middle shows schematic of F781D mutation on mGlu_1_ protomer (cyan) and T174A mutation on mGlu_5_ protomer (magenta), restricting to trans activation from mGlu_1_ to mGlu_5_. Right shows a concentration-response curve of mGlu_1/5_ WT (black), versus mGlu_1/5_ in trans from 5 to 1 (light orange), versus mGlu_1/5_ in trans from 1 to 5 (dark orange) by CODA-RET. Symbols represent the mean drug-induced BRET response and error bars represent ± SEM. The exact number of ‘n’ independent experiments and technical replicates are reported in Table S2.

### Trans activation

To assess trans activation—where glutamate binding to one protomer drives G protein coupling through the other—we co-expressed a glutamate-binding deficient protomer (mGlu_1_^T188A^ or mGlu_5_^T174A^) with a G protein coupling-deficient protomer (mGlu_1_^F781D^ or mGlu_5_^F767D^) (Figure 2A, cartoons).^8,11,24,27^ In contrast to prior work, mGlu_1/1_ homodimers showed almost no trans activity in either assay system (Figures 2A and S3A). mGlu_5/5_ homodimers retained some trans activity, though with vastly reduced potency compared to WT (Figures 2B and S3B), consistent with previous findings that both orthosteric sites must be occupied for full mGlu_5_ activation.^8,16,30,31^ In CODA-RET, trans activation of the heterodimer was impaired compared to WT; no apparent difference was observed between the two directions, but notably both achieved activity comparable to mGlu_5/5_ homodimer trans activation (Figure 2C). The GABA_B_-tail assay also showed impaired trans activation in both directions but revealed an asymmetry: trans activation through the mGlu_1_ protomer (mGlu_1_^F781S^/mGlu_5_^R68E^) yielded greater efficacy than through mGlu_5_ (mGlu_1_^R78L^/mGlu_5_^F767S^) (Figure S3C). Thus, while mGlu1 showed essentially no trans activity in the homodimer, it acquires partial trans capability in the heterodimer.

### Cis activation

We next examined cis activation—where glutamate binding and G protein coupling occur through the same protomer—by pairing a double-mutant protomer (both glutamate-binding and G protein coupling deficient) with a WT protomer (Figures 3A, 3B and 3C, cartoons). This experiment has not been reported previously for Group I mGlu receptors. The mGlu_1/1_ homodimer showed minimal cis activity in both assays (Figures 3A and S3D) and mGlu_5/5_ showed only modest cis activity in both assays, with reduced potency in CODA-RET and reduced efficacy in the GABA_B_-tail assay (Figures 3B and S3E). At the mGlu_1/5_ heterodimer, cis activation revealed a striking asymmetry in CODA-RET: signaling through the mGlu_1_ protomer preserved nearly full efficacy (∼75% of WT) but with drastically reduced potency (EC_50_ >500 µM vs 9.8 µM for WT), whereas signaling through mGlu_5_ showed reduced efficacy (∼25% of WT) but relatively better potency (EC_50_ 82 µM) (Figure 3D). The GABA_B_-tail assay confirmed this asymmetry, with cis activation through mGlu_1_ reaching full efficacy while cis activation through mGlu_5_ was reduced (Figure S3F). Thus, while an mGlu_1_ protomer in an mGlu_1/1_ homodimer exhibits relatively little cis activity, pairing with mGlu_5_ enables cis activation through mGlu_1_ to reach nearly full efficacy, a striking gain of function that emerges in the heterodimer context.

**Figure 3.**
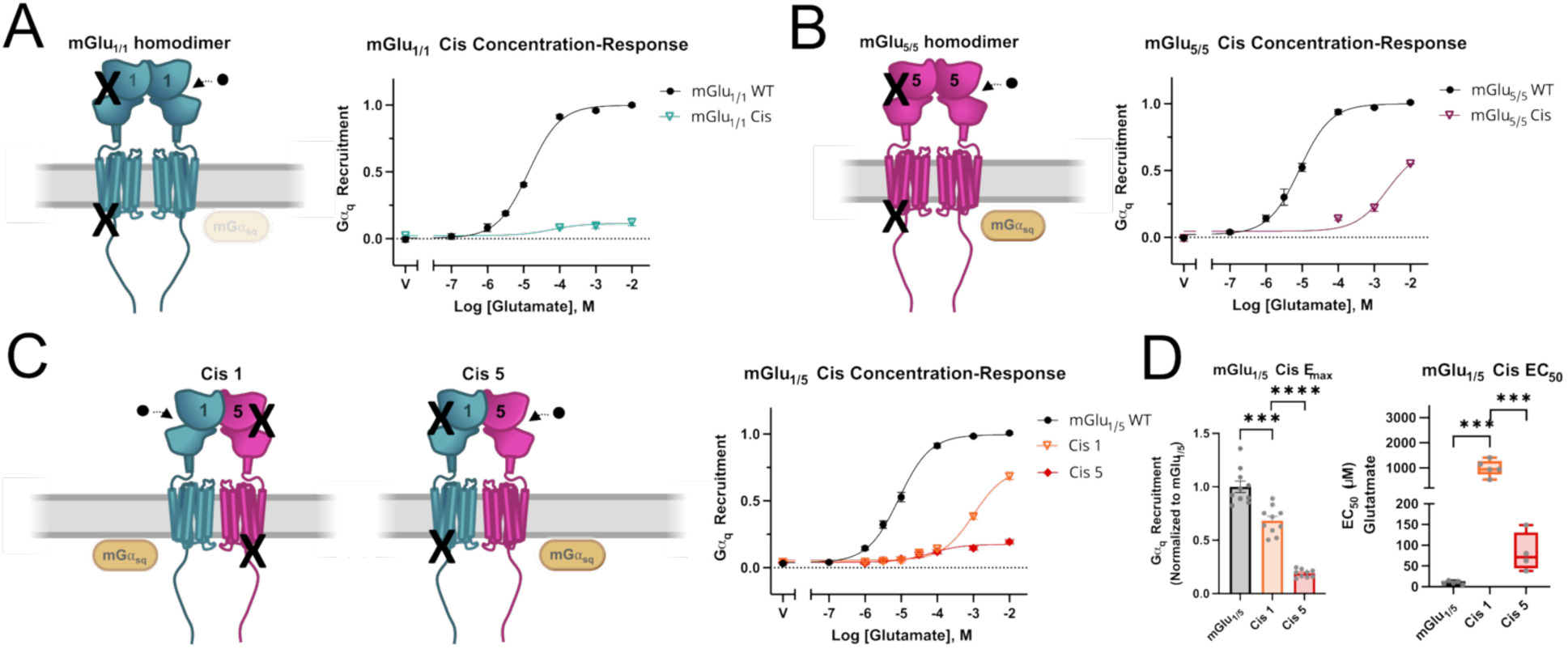
Group I mGluRs Demonstrate Poor Cis Activation and Asymmetry in the Heterodimer. (A) Left shows schematic of T188A and F781D mutations on one mGlu_1_ protomer paired with mGlu_1_ WT (cyan), restricting to cis activation. Right shows a concentration-response curve of mGlu_1/1_ WT (black) versus mGlu_1/1_ in cis (cyan) by CODA-RET. (B) Left shows schematic of T174A and F767D mutations on one mGlu_5_ protomer paired with mGlu_5_ WT (magenta), restricting to cis activation. Right shows a concentration-response curve of mGlu_5/5_ WT (black) versus mGlu_5/5_ in cis (magenta) by CODA-RET. (C) Left shows schematic of T174A and F767D mutations on mGlu_5_ protomer (magenta) paired with mGlu_1_ WT, restricting to cis activation through mGlu_1_. Middle shows T188A and F781D mutations on mGlu_1_ protomer (cyan) paired with mGlu_5_ WT (magenta), restricting to cis activation through mGlu_5_. Right shows a concentration-response curve of mGlu_1/5_ WT (black) versus mGlu_1/5_ in cis through mGlu_1_ (light orange) versus mGlu_1/5_ in cis through mGlu_5_ (dark orange) by CODA-RET. (D) Corresponding E_max_ (left) and EC_50_ (right) for Figure 3C. For E_max_: ***p = 0.0002 and ****p = < 0.0001. For EC_50,_ ***p = 0.0001, ***p = 0.0008. For plots, symbols represent the mean drug induced BRET response and error bars represent ± SEM. The exact number of ‘n’ independent experiments and technical replicates are reported in Table S2.

**Figure 4.**
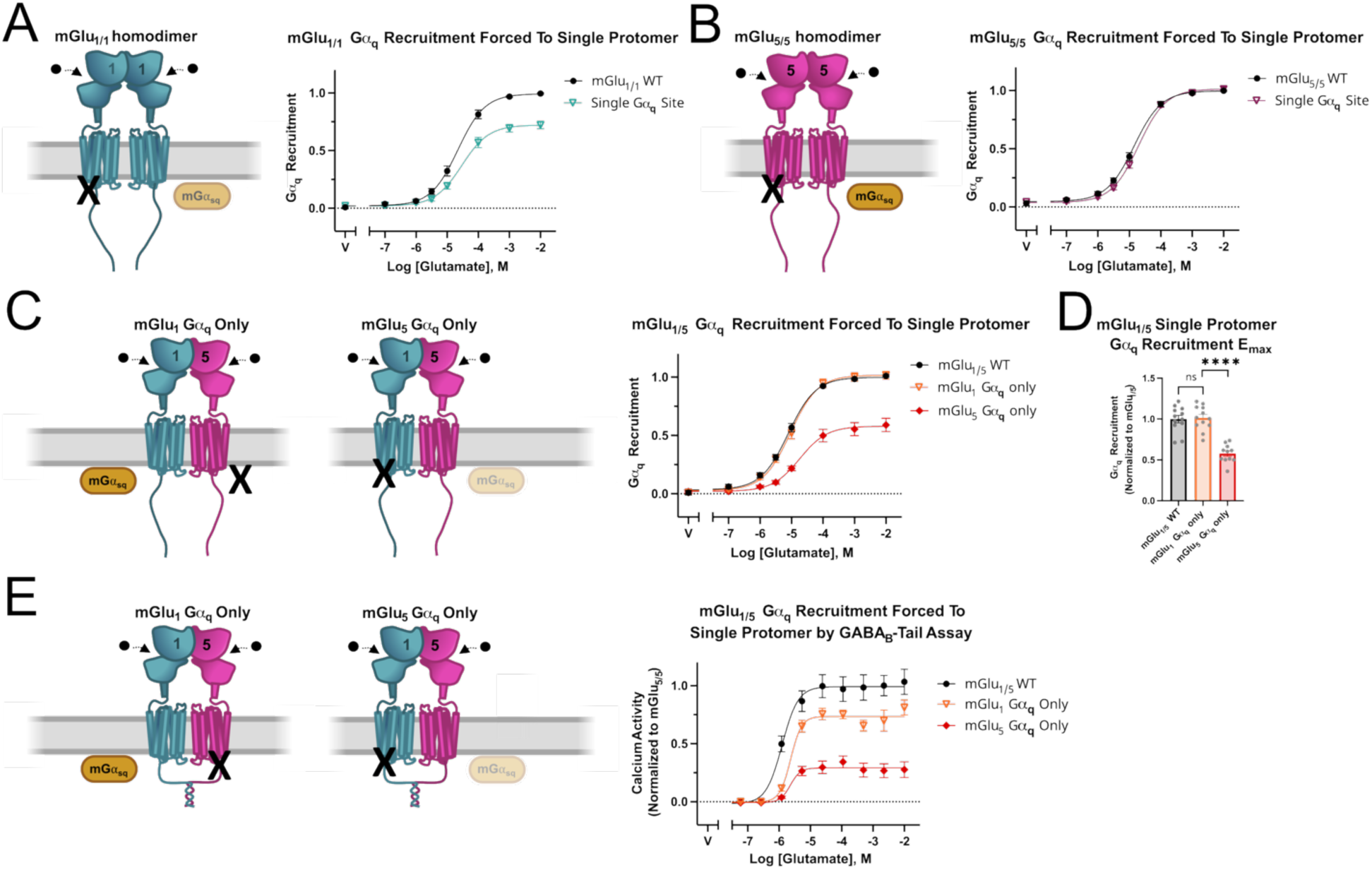
Group I Heterodimers Recruit G⍺_q_ Primarily Through mGlu_1_. (A) Left shows schematic of F781D mutation on one mGlu_1_ protomer paired with mGlu_1_ WT (cyan), restricting G⍺_q_ recruitment to one protomer. Right shows a concentration-response curve of mGlu_1/1_ WT (black) versus mGlu_1/1_ via a single protomer (cyan) by CODA-RET. (B) Left shows schematic of F767D mutation on one mGlu5 protomer paired with mGlu_5_ WT (magenta), restricting G⍺_q_ recruitment to one protomer. Right shows a concentration-response curve of mGlu_5/5_ WT (black) versus mGlu_5/5_ via a single protomer (magenta) by CODA-RET. (C) Left shows schematic of F767D mutation on mGlu5 protomer (magenta) paired with mGlu_1_ WT, restricting G⍺_q_ recruitment to mGlu_1_ protomer. Middle shows schematic of F781D mutation on mGlu_1_ protomer (cyan) paired with mGlu_5_ WT, restricting G⍺_q_ recruitment to mGlu_5_ protomer. Right shows a concentration-response curve of mGlu_1/5_ WT (black) versus mGlu_1/5_ G⍺_q_ activity via mGlu_1_ only (light orange) versus via mGlu5 only (dark orange) by CODA-RET. (D) Corresponding E_max_ for Figure 4C. ns = 0.84 and ****p = < 0.0001. (E) Left shows schematic of F767S mutation on mGlu_5_-GB2 protomer (magenta) paired with mGlu_1_-GB1 WT, restricting G⍺_q_ recruitment to mGlu_1_ protomer. Middle shows schematic of F781S mutation on mGlu_1_-GB1 protomer (cyan) paired with mGlu5-GB2 WT, restricting G⍺_q_ recruitment to mGlu_5_ protomer. Right shows a concentration-response curve of mGlu_1/5_ WT (black) versus mGlu_1/5_ G⍺_q_ activity via mGlu_1_ only (light orange) versus via mGlu5 only (dark orange) by GABA_B_-tail assay. For panels 4A-D, symbols represent the mean drug-induced BRET response and error bars represent ± SEM. For panel 4E, symbols represent the mean drug-induced calcium-sensitive fluorescence and error bars represent ± SEM. The exact number of ‘n’ independent experiments and technical replicates are reported in Table S2.

### Protomer-specific Gα_q_ recruitment

To directly assess each protomer’s contribution to Gα_q_ coupling when both protomers can bind glutamate normally, we paired a G protein coupling-deficient protomer with a WT partner, restricting Gαq recruitment to a single defined protomer(Figures 4A, 4B, and 4C, cartoons). For homodimers in CODA-RET, mGlu_1/1_ showed a small reduction in efficacy when restricted to signaling from a single protomer, while mGlu_5/5_ was essentially unaffected (Figures 4A and 4B). In marked contrast, at the mGlu_1/5_ heterodimer, the asymmetry was pronounced: when signaling was restricted to mGlu_1_ (mGlu_1_/5_F767D_), efficacy and potency were fully preserved compared to WT, but when restricted to mGlu_5_ (mGlu_1_^F781D^/ mGlu_5_), efficacy dropped substantially (Figures 4C and 4D). The GABA_B_-tail assay showed a similar pattern, with near-complete preservation of signaling through mGlu_1_ and substantially reduced signaling through mGlu_5_ (Figure 4E). These convergent results demonstrate that Gα_q_ recruitment at the mGlu_1/5_ heterodimer occurs predominantly through the mGlu_1_ protomer.

### The transmembrane domain confers mGlu_1_ signaling bias

Having established that Gα_q_ signaling at the mGlu_1/5_ heterodimer occurs predominantly through the mGlu_1_ protomer, we next asked which receptor domain is responsible for this asymmetry. mGlu receptors have a modular architecture: the extracellular domain (ECD), comprising the Venus flytrap (VFT) and cysteine-rich domain (CRD), binds glutamate and transduces conformational changes to the transmembrane domain (TM), which couples to G proteins.^1^ To determine whether the bias originates in the ECD or TM, we generated chimeric receptors swapping these domains between mGlu_1_ and mGlu_5_ (see Methods).^32–35^ For clarity, we use WT to denote the wild-type protomer in the chimeric heterodimers.

We first tested whether the mGlu_1_ ECD could confer signaling bias by pairing an mGlu_5_ TM carrying the mGlu_1_ ECD (mGlu_5_^1ECD^) with mGlu_5_^WT^. It did not, as both TM domains signaled equivalently (Figure 5A). The converse experiment, pairing the mGlu_1_ TM carrying the mGlu_5_ ECD (mGlu_1_^5ECD^) with mGlu_1_^WT^, similarly showed no bias (Figure 5B). Thus, the ECD does not determine which protomer dominates signaling.

**Figure 5.**
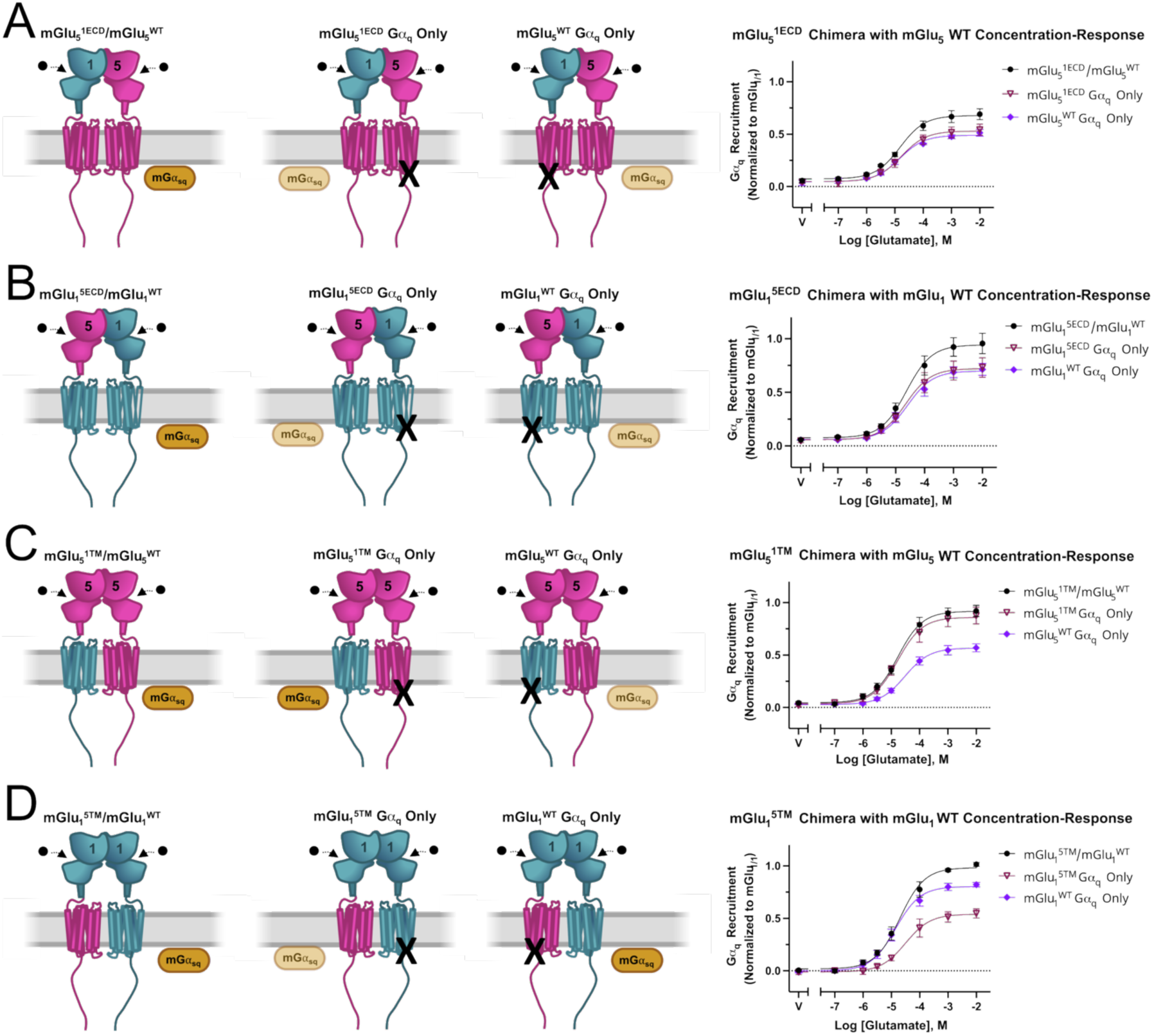
mGlu_1_ Bias Is Facilitated by the Transmembrane Domain. (A) Left shows schematics (left to right) of an mGlu_1_ ECD with mGlu_5_ TM (mGlu_5_^1ECD^) paired with mGlu_5_ WT, this same heterodimer with a F767D mutation on the mGlu_5_ WT protomer, and the heterodimer with a F767D mutation on the mGlu_5_^1ECD^ protomer. Right shows a concentration-response curve of these three heterodimers in black, burgundy, and lilac, respectively. Notably no bias signaling could be recapitulated with the mutations in these chimeric heterodimers. (B) Left shows schematics (left to right) of an mGlu_5_ ECD with mGlu_1_ TM (mGlu_1_^5ECD^) paired with mGlu_1_ WT, this same heterodimer with a F781D mutation on the mGlu_1_ WT protomer, and the heterodimer with a F781D mutation on the mGlu_1_^5ECD^ protomer. Right shows a concentration-response curve of these three heterodimers in black, burgundy, and lilac, respectively. Notably no bias signaling could be recapitulated with the mutations in these chimeric heterodimers. (C) Left shows schematics (left to right) of an mGlu_5_ ECD with an mGlu_1_ TM (mGlu_5_^1TM^) paired with mGlu_5_ WT, this same heterodimer with a F767D mutation on the mGlu_5_ WT protomer, and the heterodimer with a F781D mutation on the mGlu_5_^1TM^ protomer. Right shows a concentration-response curve of these three heterodimers in black, burgundy, and lilac respectively. Notably, bias signaling could be recapitulated with the mutations in these chimeric heterodimers. (D) Left shows schematics (left to right) of an mGlu_1_ ECD with an mGlu_5_ TM (mGlu1^5TM^) paired with mGlu_1_ WT, this same heterodimer with a F781D mutation on the mGlu_1_ WT protomer, and the heterodimer with a F767D mutation on the mGlu1^5TM^ protomer. Right shows a concentration-response curve of these three heterodimers in black, burgundy, and lilac respectively. Notably, bias signaling could be recapitulated with the mutations in these chimeric heterodimers. Symbols represent the mean drug-induced BRET response and error bars represent ± SEM. The exact number of ‘n’ independent experiments and technical replicates are reported in Table S2.

We then asked whether the TM domain itself was responsible for the bias. When we paired the mGlu_5_ ECD carrying the mGlu_1_ TM (mGlu_5_^1TM^) with mGlu_5_^WT^, signaling was biased toward the chimeric protomer containing the mGlu_1_ TM (Figure 5C). The reciprocal experiment confirmed this: pairing the mGlu_1_ ECD carrying the mGlu_5_ TM (mGlu_1_^5TM^) with mGlu_1_^WT^ shifted bias toward the mGlu_1_^WT^—away from the protomer carrying the mGlu_5_ TM (Figure 5D). In both cases, the mGlu_1_ TM dominated signaling regardless of which ECD was attached to it.

These experiments demonstrate that when mGlu_1_ and mGlu_5_ TM domains are present in the same dimer, the mGlu_1_ TM preferentially couples to Gα_q_. This property is autonomous—it does not depend on the ECD—and provides a functional explanation for why Gα_q_ signaling in the mGlu_1/5_ heterodimer flows predominantly through the mGlu_1_ protomer. Because allosteric modulators bind within the TM domain, this finding has direct implications for how PAMs and NAMs might behave at the heterodimer.

### Allosteric modulator effects at the mGlu_1/5_ heterodimer

Given that Gα_q_ signaling at the mGlu_1/5_ heterodimer flows predominantly through the mGlu_1_ TM, we predicted that allosteric modulators—which bind within the TM—might behave differently at the heterodimer than at homodimers. We tested this using the mGlu_1_-specific positive allosteric modulator (PAM) VU6024578 and negative allosteric modulator (NAM) FITM, as well as the mGlu_5_-specific NAM, MTEP.^36,37^

### mGlu_1_ PAM and NAM

As expected, VU6024578 (PAM) enhanced and FITM (NAM) inhibited signaling at the mGlu_1/1_ homodimer, while neither compound affected the mGlu_5/5_ homodimer, confirming their selectivity (Figure 6A and 6B). At the mGlu_1/5_ heterodimer, both compounds retained their expected activity: the PAM enhanced and the NAM inhibited signaling (Figures 6C and S4A).

**Figure 6.**
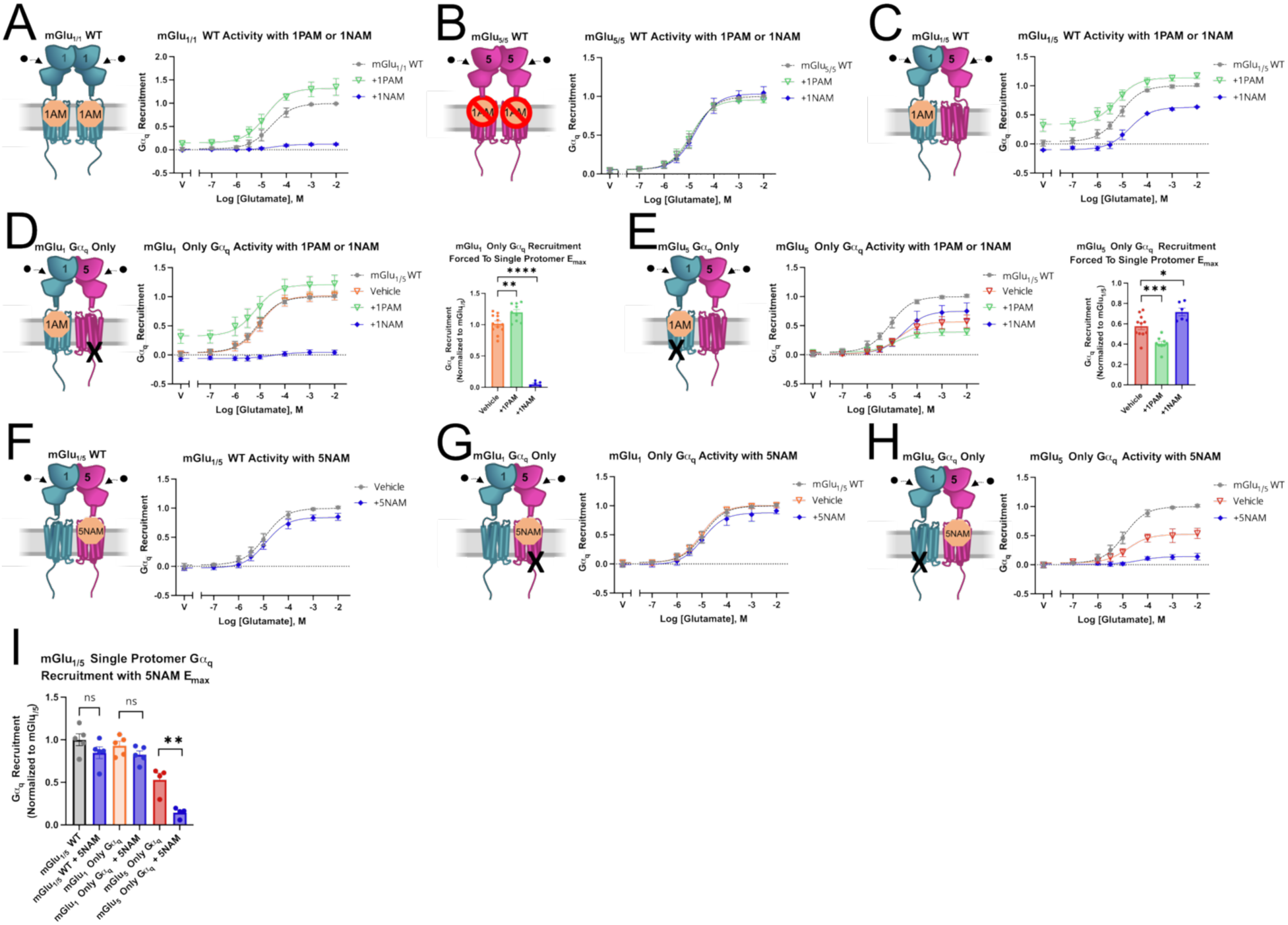
Inversion of mGlu_1_ PAM and NAM Signaling and Limited Effects of the mGlu5 NAM. (A) Left shows a schematic of mGlu_1/1_ WT with a mGlu_1_ PAM (1PAM) bound to the allosteric binding pocket. Right shows a concentration-response curve of mGlu_1/1_ WT with vehicle (grey), with PAM (100 nM VU6024578-green), or with NAM (100 nM FITM-blue). (B) Left shows a schematic of mGlu_5/5_ WT with a 1PAM unable to bind to the allosteric binding pocket. Right shows a concentration-response curve of mGlu_5/5_ WT with vehicle (grey), with PAM (100 nM VU6024578-green), or with NAM (100 nM FITM-blue). (C) Left shows a schematic of mGlu_1/5_ WT with a 1PAM bound to the allosteric binding pocket of mGlu_1_. Right shows a concentration-response curve of mGlu_1/5_ WT with vehicle (grey), with PAM (100 nM VU6024578-green), or with NAM (100 nM FITM-blue). (D) Left shows a schematic of mGlu_1/5_ heterodimer with a G⍺_q_-binding point mutation (F767D) in mGlu_5_ with a 1PAM bound to the allosteric binding pocket of mGlu_1_. Middle shows a concentration-response curve of mGlu_1/5_ WT with vehicle (grey), mGlu_1_^WT^/mGlu_5_^F767D^ with vehicle (orange), mGlu_1_^WT^/mGlu_5_^F767D^ with 1PAM (100 nM VU6024578-green), or mGlu_1_^WT^/mGlu_5_^F767D^ with NAM (100 nM FITM-blue). Right shows corresponding E_max_: **p = 0.01 and ****p = < 0.0001. (E) Left shows a schematic of mGlu_1/5_ heterodimer with a G⍺_q_-binding point mutation (F781D) in mGlu_1_ with a 1PAM bound to the allosteric binding pocket of mGlu_1_. Middle shows a concentration-response curve of mGlu_1/5_ WT (grey) with vehicle, mGlu_1_^F781D^/mGlu_5_^WT^ with vehicle (orange), mGlu_1_^F781D^/mGlu_5_^WT^ with 1PAM (100 nM VU6024578-green), or mGlu_1_^F781D^/mGlu_5_^WT^ with 1NAM (100 nM FITM-blue). Right shows corresponding E_max_: *p = 0.02 and ***p = 0.0007. (F) Left shows a schematic of mGlu_1/5_ WT with a 5NAM bound to the allosteric binding pocket of mGlu_5_. Right shows a concentration-response curve of mGlu_1/5_ WT with vehicle (grey), with 5NAM (1uM MTEP-blue). (G) Left shows a schematic of mGlu_1/5_ heterodimer with a G⍺_q_-binding point mutation (F767D) in mGlu_5_ with a 5NAM bound to the allosteric binding pocket of mGlu_5_. Right shows a concentration-response curve of mGlu_1/5_ WT with vehicle (grey), mGlu1^WT^/mGlu5^F767D^ with vehicle (orange), and mGlu_1_^WT^/mGlu_5_^F767D^ with 5NAM (1 uM MTEP-blue). (H) Left shows a schematic of mGlu_1/5_ heterodimer with a G⍺_q_-binding point mutation (F781D) in mGlu_1_ with a 5NAM bound to the allosteric binding pocket of mGlu_5_. Right shows a concentration-response curve of mGlu_1/5_ WT with vehicle (grey), mGlu_1_^F781D^/mGlu_5_^WT^ with vehicle (orange), and mGlu_1_^F781D^/mGlu_5_^WT^ with 5NAM (1 uM MTEP-blue). (I) Corresponding E_max_ for Figures 6F-6H. ns = 0.16, ns = 0.16 and **p = 0.004. Symbols represent the mean drug-induced BRET response and error bars represent ± SEM. The exact number of ‘n’ independent experiments and technical replicates are reported in Table S2.

We then tested how these modulators behaved when signaling was restricted to a single protomer. When Gα_q_ recruitment was restricted to mGlu_1_ (mGlu_1_/mGlu_5_^F767D^), the PAM and NAM produced effects nearly identical to those at the mGlu_1/1_ homodimer (Figures 6D and S4B). However, when Gα_q_ recruitment was restricted to mGlu_5_ (mGlu_1_^F781D^/mGlu_5_), the effects inverted: remarkably, the mGlu_1_ “PAM” now reduced signaling, and the mGlu_1_ “NAM” increased it (Figures 6E and S4C). Thus, simply by changing which protomer could couple to Gα_q_, a positive modulator became inhibitory and a negative modulator became stimulatory. This inversion suggests that the dominant mGlu_1_ TM suppresses mGlu_5_ signaling within the heterodimer, even when mGlu_1_ cannot couple Gα_q_ directly. Stabilizing the inactive mGlu_1_ conformation with a NAM relieves this suppression.

### mGlu_5_ NAM

To further probe mGlu_5_’s role in heterodimer signaling, we tested the mGlu_5_-selective NAM MTEP. At the WT heterodimer, MTEP had only a small effect on signaling (Figure 6F)—consistent with signaling flowing predominantly through mGlu_1_, rendering mGlu_5_ inhibition largely inconsequential. This small effect persisted even when mGlu_5_ could not couple to Gα_q_ (mGlu_1_/mGlu_5_^F767D^; Figure 6G), consistent with a minor trans inhibition of mGlu_1_ rather than direct suppression of mGlu_5_ signaling. However, when signaling was restricted to mGlu_5_ (mGlu_1_^F781D^/mGlu_5_), MTEP dramatically reduced the efficacy, confirming that mGlu_5_ retains full sensitivity to cis inhibition when restricted to serve as the Gα_q_-coupling protomer (Figures 6H and 6I).

### mGlu_1_ PAM VU6024578 and NAM FITM act in cis within the dimer

These heterodimer-specific modulator effects raised the question of whether these effects emerge specifically from the heterodimer context or reflect underlying mechanisms that are latent in homodimers but undetectable because both protomers are identical and equally capable of coupling to Gα_q_. To determine whether modulator binding to one protomer affects signaling through that same protomer or through its partner at the mGlu_1/1_ homodimer, we introduced mutations in the allosteric binding pocket that prevent modulator binding while preserving receptor function.

For the PAM VU6024578, we introduced the mutation Y672V based on an analogous mutation in mGlu_5_ (Y658V) from previous work.^16,38,39^ The mutation abolished PAM activity of VU6024578 at the mGlu_1/1_ homodimer (Figures 7A and 7B). We then combined this mutation with the Gα_q_-coupling disrupting mutation (F781D) in defined configurations. When both mutations were placed on the same protomer (mGlu_1_^Y672V/F781D^ paired with mGlu_1_), the PAM retained full activity (Figure 7C)—it could still bind to and potentiate the WT protomer in cis configuration. However, when the PAM-disrupting mutation and Gα_q_-coupling mutation were placed on opposite protomers (mGlu_1_^Y672V^ paired with mGlu_1_^F781D^), the PAM had no effect (Figure 7D), consistent with an inability of the PAM to act in trans. In this configuration, the PAM can only bind the protomer that cannot couple to Gα_q_, so the absence of any effect demonstrates that the PAM can only potentiate Gα_q_ coupling through the protomer to which it is bound.

**Figure 7.**
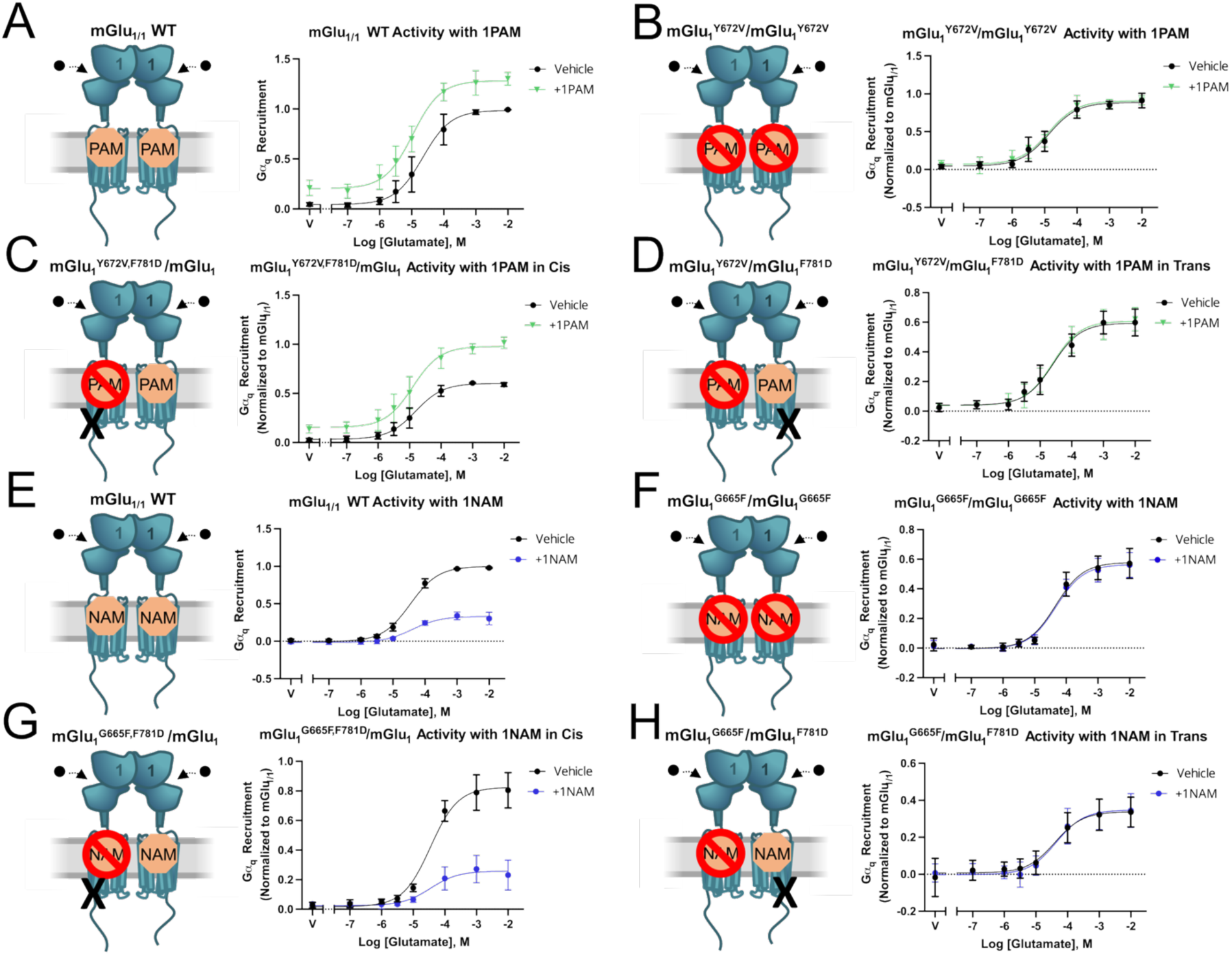
mGlu_1_ 1PAM and 1NAM Signaling In Cis. (A) Left shows a schematic of mGlu_1/1_ WT with a 1PAM able to bind to either allosteric binding pocket. Right shows a concentration-response curve of mGlu_1/1_ WT with vehicle (black) or with 1PAM (100 nM VU6024578-green). (B) Left shows a schematic of mGlu_1_^Y672V^/mGlu_1_^Y672V^ with a 1PAM unable to bind to the allosteric binding pocket and/or produce effect. Right shows a concentration-response curve of mGlu_1_^Y672V^/ mGlu1^Y672V^ with vehicle (black) or with PAM (100 nM VU6024578-green). Activity is relative to mGlu_1/1_ WT. (C) Left shows a schematic of mGlu_1_^Y672V,F781D^/mGlu_1_ with a 1PAM able to bind the allosteric binding pocket on the same protomer that can bind G⍺_q_ (acting in cis). Right shows a concentration-response curve of mGlu_1_^Y672V,F781D^/mGlu_1_ with vehicle (black) or with PAM (100 nM VU6024578-green). Activity is relative to mGlu_1/1_ WT. (D) Left shows a schematic of mGlu_1_^Y672V^/mGlu_1_^F781D^ with a 1PAM able to bind the allosteric binding pocket on the adjacent protomer that can bind G⍺_q_ (acting in trans). Right shows a concentration-response curve of mGlu_1_^Y672V^/mGlu_1_^F781D^ with vehicle (black) or with 1PAM (100 nM VU6024578-green). Activity is relative to mGlu_1/1_ WT. (E) Left shows a schematic of mGlu_1/1_ WT with a 1NAM able to bind to either allosteric binding pocket. Right shows a concentration-response curve of mGlu_1/1_ WT with vehicle (black) or with NAM (100 nM FITM-blue). (F) Left shows a schematic of mGlu_1_^G665F^/mGlu_1_^G665F^ with a 1NAM unable to bind to the allosteric binding pocket and/or produce effect. Right shows a concentration-response curve of mGlu_1_^G665F^/mGlu_1_^G665F^ with vehicle (black) or with NAM (100 nM FITM-blue). Activity is relative to mGlu_1/1_ WT. (G) Left shows a schematic of mGlu_1_^G665F,F781D^/mGlu_1_ with a 1NAM able to bind the allosteric binding pocket on the same protomer that can bind G⍺_q_ (acting in cis). Right shows a concentration-response curve of mGlu_1_^G665F,F781D^/mGlu_1_ with vehicle (black) or with 1NAM (100 nM FITM-blue). Activity is relative to mGlu_1/1_ WT. (H) Left shows a schematic of mGlu_1_^G665F^/mGlu_1_^F781D^ with a 1NAM able to bind the allosteric binding pocket on the adjacent protomer that can bind G⍺_q_ (acting in trans). Right shows a concentration-response curve of mGlu_1_^G665F^/mGlu_1_^F781D^ with vehicle (black) or with 1NAM (100 nM FITM-blue). Activity is relative to mGlu_1 /1_ WT. Symbols represent the mean drug-induced BRET response and error bars represent ± SEM. The exact number of ‘n’ independent experiments and technical replicates are reported in Table S2.

For the NAM FITM, we introduced the mutation G665F based on analogous mutations in mGlu_5_ from previous work.^39^ The mutation abolished NAM activity of FITM at the mGlu_1/1_ homodimer (Figures 7E and 7F). The same logic applied: when the NAM-disrupting mutation and Gα_q_-coupling mutation were on the same protomer (mGlu_1_^G665F/F781D^ paired with mGlu_1_), the NAM retained activity (Figure 7G). When on opposite protomers (mGlu_1_^G665F^ paired with mGlu_1_^F781D^), the NAM had no effect (Figure 7H). Thus, like the PAM, the NAM can only inhibit Gα_q_ coupling through the protomer to which it is bound.

## Discussion

The prevailing view of mGlu_1/5_ heterodimers held that both protomers contribute symmetrically to signal transduction.^11^ Using CODA-RET to measure direct Gα_q_ recruitment to intact, full-length receptors, we find instead that signaling is markedly asymmetric: the mGlu_1_ protomer dominates. When Gα_q_ coupling was restricted to mGlu_5_, efficacy dropped substantially; when restricted to mGlu_1_, signaling was fully preserved. This asymmetry was consistent across trans activation, cis activation, and protomer-specific Gα_q_ recruitment experiments, and was confirmed with orthogonal GABA_B_-tail calcium mobilization assays, in which a linker modification potentially relieved steric constraints present in the previously published constructs.

Beyond asymmetry, we observed a striking gain of function. As homodimers, mGlu_1/1_ exhibited almost no trans or cis activity, consistent with the requirement that both VFTs close for full activation, whereas mGlu_5/5_ showed partial activity in both modes.^8,14,15,24^ Yet in the heterodimer, mGlu_1_ acquired trans and cis activity comparable to what mGlu_5/5_ achieves as a homodimer, and, notably, cis activation through mGlu_1_ preserved efficacy while cis activation through mGlu_5_ preserved potency, suggesting each protomer contributes distinct functional properties to the complex. Pairing with mGlu_5_ enabled mGlu_1_ to function in ways it did not when paired with itself, suggesting that mGlu_5_, despite contributing minimally to Gα_q_ coupling itself, enables mGlu_1_ to adopt conformations or engage in dimer rearrangements that it cannot achieve when paired with another mGlu_1_. The heterodimer is thus not simply an intermediate between two homodimers but a receptor complex with emergent signaling properties— one in which pairing fundamentally reshapes the signaling capacity of each protomer.

Chimeric receptor experiments localized the structural basis of mGlu_1_ dominance to the transmembrane domain. Swapping ECDs between mGlu_1_ and mGlu_5_ did not transfer the bias, whereas swapping TM domains did. In every configuration, the mGlu_1_ TM preferentially coupled to Gα_q_ regardless of which ECD was attached. This identifies the TM as an intrinsically more effective Gα_q_-coupling module when mGlu_1_ and mGlu_5_ TM domains are present in the same dimer. The structural features that confer this advantage remain to be determined.

The dominance of the mGlu_1_ TM has pharmacological consequences. At the WT heterodimer, where signaling flows predominantly through mGlu_1_, VU6024578 and FITM retained their expected activities. However, when signaling was restricted to mGlu_5_, the “PAM” became inhibitory and the “NAM” became stimulatory, a phenomenon we term allosteric inversion. This inversion suggests that the mGlu_1_ TM suppresses mGlu_5_ signaling within the heterodimer, and that stabilizing an inactive mGlu_1_ conformation with a NAM relieves this suppression.

Conversely, the mGlu_5_ NAM MTEP had minimal direct effect at the heterodimer, consistent with the limited contribution of mGlu_5_ to overall signaling. Using allosteric binding-site mutations, we demonstrated that VU6024578 and FITM act exclusively in cis, modulating only the protomer to which they bind. At homodimers, cis versus trans activity is pharmacologically equivalent as both protomers are identical. At heterodimers, however, the distinction becomes critical: a cis-acting modulator’s effect depends entirely on the signaling capacity of the protomer it occupies.

These findings have implications for therapeutic targeting of Group I mGlu receptors. Because mGlu_1_ and mGlu_5_ are co-expressed in many brain regions and can form heterodimers, drugs developed against homodimers may behave differently in circuits where heterodimers predominate. Since cis-acting modulators like VU6024578 depend entirely on the signaling capacity of the protomer they occupy, at the mGlu_1/5_ heterodimer, their effects flow through mGlu_1_. Whether trans-acting modulators exist, and how they might behave at heterodimers, remains an important question. Modulators that potentiate signaling across protomers could in principle engage the mGlu_5_ protomer more effectively, or conversely, might lose activity altogether in the heterodimer if they drive signaling through a protomer with limited Gα_q_-coupling capacity. By extension, compounds that selectively modulate mGlu_1/1_ homodimers while sparing—or differentially affecting— mGlu_1/5_ heterodimers could have distinct therapeutic profiles, a possibility that warrants further investigation.

In summary, we have shown that the mGlu_1/5_ heterodimer is not a symmetric receptor complex but one in which Gα_q_ signaling flows almost exclusively through mGlu_1_. This asymmetry is conferred by the transmembrane domain and has profound consequences for allosteric pharmacology. These findings suggest that homodimer-versus heterodimer-selective pharmacology may be achievable and therapeutically relevant. Understanding these properties will be essential for developing drugs that effectively target Group I mGlu receptors in the brain.

### Limitations of the Study

Several limitations bear noting. Our experiments were conducted in heterologous cells, though the consistency across two independent assay systems (CODA-RET and GABA_B_-tail calcium mobilization) and the agreement with known pharmacology of the homodimers support the validity of our findings. Confirmation in native neurons at physiological expression levels will be an important next step. While Group I mGlu receptors signal predominantly through Gα_q_, whether protomer contributions differ for other signaling pathways remains unexplored. Finally, the abundance and regulation of mGlu_1/5_ heterodimers in specific brain circuits requires further investigation.

## Acknowledgements

This work was supported by National Institutes of Health grants R01 NS132060 (J.A.J.), R01 MH054137 (J.A.J.), the Hope for Depression Research Foundation (J.A.J.), St. Jude Children’s Research Hospital GPCR Collaborative (J.A.J.), R01 MH062646 (P. Jeffrey Conn), R01 NS031373 (P. Jeffrey Conn), R01 MH119673 (P. Jeffrey Conn), R25 MH125775-01 (Jeremy Veenstra-Vanderweele), R25 MH086466 (Melissa Arbuckle), and the Leon Levy Foundation (J.B.S.). This work was also supported by the American Academy of Child and Adolescent Psychiatry (AACAP) Pilot Research Award for Junior Faculty and Child and Adolescent Psychiatry Fellows supported by AACAP (J.B.S.) and Drs. Gabrielle and Harold Carlson Psychopharmacology Research Award (J.B.S.); its contents are the responsibility of the authors and do not necessarily reflect the official views of AACAP. We also thank Nevin Lambert for providing miniG plasmids. We thank Jason Manka for the synthesis of FITM and the Warren family for endowing the Warren Center for Neuroscience Drug Discovery.

## Author Contributions

J.B.S., C.N., and J.A.J. designed the live-cell assays and interpreted the results with critical input from W.B.A. and X. Lin. J.B.S. and M.L. performed the live-cell CODA-RET and GCaMP assays. X. Lei performed the GABA_B_-tail assays. P.S. synthesized VU6024578. J.B.S., X.L., M.L., X. Lin, and A.R. performed data analysis. J.B.S. and J.A.J. wrote the manuscript, with contributions from all the authors.

## Methods

### Chimeras

mGlu_1_/mGlu_5_ chimeras were generated by separating the Venus-fly-trap/cysteine-rich domains from the transmembrane domain at junctional sites P581 for mGlu_1_ and P568 for mGlu_5_ based on previous work with mutagenesis and calcium-sensing receptor (CaR) chimeras.^32,34^ Reference Table S1 for full construct details.

### Reagents and Ligands

FITM (4-fluoro-*N*-[4-[6-(isopropylamino)pyrimidin-4-yl]-1,3-thiazol-2-yl]-*N*-methylbenzamide) and MTEP (3-((2-Methyl-1,3-thiazol-4-yl)ethynyl)pyridine hydrochloride) were purchased from Tocris Bioscience (Bristol, UK). VU6024578 was synthesized at Vanderbilt University.^36^ 10 mM stock solutions of FITM, MTEP, and VU6024578 were prepared by dissolving them in dimethyl sulfoxide (DMSO) (Sigma-Aldrich, St. Louis, MO). 1 M L-glutamate (Sigma-Aldrich, St. Louis, MO) was made in PBS, using Sodium Hydroxide solution (10.0 N) (Fisher Scientific, Waltham, MA) to bring pH to 7.0. Polyethylenimine Hydrochloride (PEI Max) was purchased from Kyfora Bio (Horsham, PA) and dissolved in sterile water at 1 mg/mL. Furimazine was purchased from BenchChem (Austin, TX) and dissolved in 90% ethanol 10% glycerol to a 5 mM stock solution.

### Plasmids

For CODA-RET and GCaMP assays, pcDNA3.1 (+) expression plasmids containing full-length rat mGlu_1_ and mGlu_5_ protomers with HiBiT and LgBiT were constructed as described in the Results section and Table S1 using NEBuilder® HIFI DNA Assembly Cloning Kit. The mVenus-miniG_sq_ plasmid was courtesy of the Nevin Lambert Lab and the EAAT1 plasmid was courtesy of the Joshua Levitz Lab.^40^ Point mutations were generated using either Q5® Site-Directed Mutagenesis or NEBuilder® HIFI DNA Assembly Cloning Kit (New England Biolabs, Ipswich, MA). DNA sequence was confirmed by Genewiz (Chelmsford, MA) with Sanger sequencing and Plasmidsaurus (South San Francisco, CA) whole plasmid sequencing.

For GABA_B_-tail assays, mGlu_1_ and mGlu_5_ receptor plasmids were constructed with GB1 and GB2 tails as described in the following publications and full sequences are shown in Table S1.^11,14,41^ A hemagglutinin (HA) tag sequence was inserted after the signal peptide in the mGlu_1_ N-terminus. The rat mGlu_1_ C-terminus was replaced after H859 with three alanine residues, followed by the sequence of the GB1 or GB2 tail, and then a KKTN endoplasmic reticulum retention signal. The entire sequence was fused into pcDNA3 via BamHI and XhoI restriction sites. A FLAG-tag sequence was inserted after the signal peptide in the mGlu_5_ N-terminus. The rat mGlu_5_ C-terminus was replaced after H855 with three alanine residues, followed by the sequence of the GB1 or GB2 tail, and then a KKTN endoplasmic reticulum retention signal. The entire sequence was fused into pcDNA3 via BamHI and XbaI restriction sites. Mutations (R78L, F781S for mGlu_1_ and R68E, F767S for mGlu_5_) were generated using a Q5® Site-Directed Mutagenesis Kit (E0554 NEB) with designed primers. DNA sequence was confirmed by Genewiz (Chelmsford, MA) with Plasmid-EZ sequencing.

### Cell culture conditions

For CODA-RET cell-based assays, HEK293T cells derived from embryonic kidney cells, purchased from Sigma, were used. Cells were maintained in 10 mL of Dulbecco’s modified Eagle’s medium (DMEM), high glucose (Gibco, Waltham, MA), supplemented with 100 Units/mL penicillin-streptomycin (Corning Inc., Corning, NY) and 10% fetal bovine serum (FBS) (Corning Inc., Corning, NY) at 37 °C under 5% CO2. The cells were passaged using 0.05% trypsin-EDTA phenol red (Thermo-Fisher, Waltham, MA) at ∼80% confluency.

For GABA_B_-tail cell-based assays, HEK293A cells were maintained in growth medium containing 90% Dulbecco’s Modified Eagle Media (DMEM), 10% fetal bovine serum (FBS), 100 Units/mL penicillin/streptomycin, 20 mM HEPES, 1 mM sodium pyruvate, 2 mM L-glutamine, 1x non-essential amino acids. All cell culture reagents were purchased from Invitrogen (Carlsbad, CA).

### Transfection

For CODA-RET assays, 48 hours before transfection, 1×10^6^ HEK cells were plated in 10 cm tissue culture dishes with 10 mL DMEM and grown to ∼80% confluency at 37 °C under 5% CO_2_. 30 minutes prior to transfection, the growth medium was replaced with 8 mL of fresh DMEM and placed back in an incubator to equilibrate.

For the CODA-RET and GCaMP assays, plasmids encoding wild type and/or mutant mGlu_1_ or mGlu_5_ protomers (mGlu_1/1_ homodimer expression: 50 ng mGlu_1_-HiBit and 50 ng mGlu_1_-LgBit; mGlu_5/5_ homodimer expression: 100 ng mGlu_5_-HiBit and 100 ng mGlu_5_-LgBit; mGlu_1/5_* heterodimer: 50 ng mGlu_1_-HiBit and 100 ng mGlu_5_-LgBit; mGlu_1/5_ heterodimer: 100 ng mGlu_5_-LgBit and 50 ng mGlu_1_-HiBit) were co-transfected with 2000 ng mGsq-mVenus(+) and 1600 ng hEAAT1 into HEK293T cells using a PEI MAX ratio of 9:1. PEI MAX is mixed with Opti-MEM, reduced serum medium, GlutaMAX Supplement (Gibco, Waltham, MA) and 500 µL is added to 500 µL of Opti-MEM and DNA. This mixture is incubated at room temperature for 20 minutes and then added dropwise to each corresponding 10 cm plate. CODA-RET experiments with the heterodimer in the main figures use mGlu_1/5_ (mGlu_5_-LgBit and mGlu_1_-HiBit) but polarity controls with mGlu_1/5_* (mGlu_1_-HiBit and mGlu_5_-LgBit) for critical findings were done and found in the Supplemental Information.

For GABA_B_-tail assays, transfections were performed according to a Lipofectamine 3000 protocol (Invitrogen, Waltham, MA) following the manufacturer’s instructions. Lipofectamine 3000 reagent and DNA (2 µg for single GB tailed plasmid, 4 µg total) were diluted in Opti-MEM medium separately and then combined with P3000 to form a DNA-lipid complex. After a 15-minute incubation at room temperature, the complexes were applied dropwise to HEK 293A cells in a 10 cm dish with Opti-MEM. The lipofectamine/DNA mixtures incubated with the cells for 24 hours, after which the cells were seeded 20,000 cells/20 µL/well into Assay Medium (DMEM containing 10% dialyzed FBS, 20 mM HEPES, 100 units/mL penicillin/streptomycin, and 1 mM sodium pyruvate) on a poly-D-lysine coated 384 black well plate (Corning BioCoat) for another 24 hours before testing.

### CODA-RET assay

Following expression, the growth media was removed and the cells were washed with 5 mL of room temperature dPBS. The cells were then dissociated from their 10 cm plate using 4 mL Enzyme-free cell-dissociation solution, PBS-Based (1X) (Millipore, Burlington, MA). The cells were then diluted with enough DMEM to obtain a concentration of 5×10^5 cells/mL; for plates at ∼80% confluency this typically required ∼20 mL DMEM. Using a RAININ E4 XLS multi-pipetter (E12-300XLS+) with 300 µL LTS compatible tips for RAININ pipettes, low ret, 100 µL of cells were distributed in black frame, white well PDL coated 96-well plates (Revvity, Waltham, MA), to get to ∼5×10^4 cells/ well. The cells were then placed in the incubator for at least four hours. Then the original growth media was removed, cells were washed in 100 µL of room temperature dPBS and replaced with 70 μL of DMEM, high glucose, no glutamine, no phenol red (Gibco, Waltham, MA) supplemented with 25 mM HEPES (7.4 pH) (Gibco, Waltham, MA), 1% penicillin-streptomycin, and 1% GlutaMax (Gibco, Waltham, MA) to be starved overnight. After overnight starvation, 10 µL of 5 mM furimazine is added to each well and the baseline fluorescence and luminescence signals were quantified using the BMG Labtech Lumistar Omega. Following two reads, the machine was paused and 20 µL of the indicated glutamate, NAM, PAM or vehicle in DMEM MAX were added to each well. The BRET ratio was determined by calculating the ratio of mVenus signal (535 nm) divided by the NanoLuc (475 nm) signal.^42^ The results are expressed as the BRET change produced by the corresponding ligands.

### GCaMP assay

Following expression, the cells were dissociated, washed and plated as described in the above CODA-RET assay. The cells were then placed in the incubator overnight, and the original media was removed and cells were washed with 100 µL of room temperature dPBS and replaced with 80 µL dPBS(+) (dPBS, 0.9 mM CaCl_2_, 0.5 mM MgCl_2_, 1 g/L glucose) supplemented with 25 mM HEPES to be starved for an hour. After an hour of starvation, the baseline fluorescence and luminescence signals were quantified using a Pherastar FS plate reader (BMG Labtech). Following two reads, the machine was paused and 20 µL of the indicated glutamate, NAM, PAM or vehicle in dPBS(+) were added to each well. The BRET ratio was determined by calculating the ratio of mVenus signal (535 nm) divided by the NanoLuc (475 nm) signal.^42^ The results were expressed as the BRET change produced by the corresponding ligands.

### GABA_B_-tail Calcium Mobilization Assays

The mGlu dimers were transiently transfected into HEK293A cells as described above. 20,000 cells/20 µL/well were plated in black-walled, clear-bottomed, poly-D-lysine 384-well assay plates (Corning BioCoat) with Assay Medium. The cells were grown at 37 °C, 5% CO_2_ overnight. The medium was removed, and cells were washed with Assay Buffer (Hank’s balanced salt solution, 20 mM HEPES and 2.5 mM Probenecid (Sigma-Aldrich, St. Louis, MO)) leaving 20 μL. Twenty μL of 2.3 μM Fluo-4(2x), AM (Life Technologies, Grand Island, NY) prepared as a 2.3 mM stock in DMSO and mixed in a 1:1 ratio with 10% (w/v) pluronic acid F-127 was diluted in Assay Buffer and applied to cells for 45 minutes at 37 °C in the presence of 5% CO_2_. Dye was washed with Assay Buffer three times, leaving 20 μL of Assay Buffer in each well. Calcium flux was measured using the FDSS µCell (Functional Drug Screening System µCell, Hamamatsu, Japan), first establishing a fluorescence baseline by recording 2 images at 1 Hz (excitation, 480 ± 20 nm; emission, 540 ± 30 nm). 20 μL vehicle (DMSO) or a constant concentration (2x) of modulator solutions was then added to the cells. After 144 seconds, 10 μL of a serial dilution of glutamate (5x) was added to the cells, and the response of the cells was measured for another 156 seconds.

### Plotting and statistics

The data collected from CODA-RET and GCaMP live cell assays were plotted and fit using GraphPad Prism 10 (GraphPad Software, La Jolla, CA). Concentration-response curves of agonist, PAM, and NAM responses were analyzed by nonlinear regression to the log(agonist) vs. response model with a standard Hill Slope equal to 1 and bottom fit constrained to 0 with at least 3 independent experiments, and error bars are standard error of the mean (SEM). For the determination of *p* values, we used an unpaired two-sided t-test, where comparison with *p* < 0.05 were considered statistically significant. The technical replicates and exact number of ‘*n*’ independent experiments can be seen in Table S2. The pEC_50_ and E_max_ distributions determined from select concentration-response curves and statistical comparisons were shown in Table S2. The baseline luminescence signals from the complemented NLuc for each mGlu homodimer and heterodimer combination tested in our CODA-RET assay were similar in magnitude and shown in Figure S2.

For GABA_B_-tail assays, calcium fluorescence was recorded as fold over basal fluorescence. The raw data were analyzed by first obtaining the peak value of the calcium flux curve then subtracting the fluorescence base point obtained from vehicle and buffer added for each cell type. The final data were normalized to the maximal response to glutamate (% Glu Max) with mGlu_5_ GB1/GB2 in the presence of vehicle. Potency (EC_50_) and maximum response for compounds were determined using a four-parameter logistical equation in GraphPad Prism (La Jolla, CA):

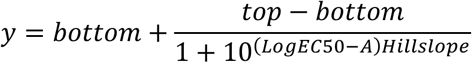

where *A* is the molar concentration of the compound; *bottom* and *top* denote the lower and upper plateaus of the concentration-response curve; HillSlope is the Hill coefficient that describes the steepness of the curve; and EC_50_ is the molar concentration of compound required to generate a response halfway between the *top* and *bottom*.

