## Supplemental Information for "Transmembrane Domain Dominance Drives Emergent Signaling and Allosteric Inversion in mGlu_1/5_ Heterodimers"

| Construct | Primary Structure |
|---|---|
| ALFA-rmGlu$_1$ WT | MVLLLILSVLLLKEDVRGSAQSTRPSRLEEELRRRLTEPDKLASSQRSVARMDGDVIIGALFSVHHQPPAEKVPERKCGEIREQYGIQRVEAMFHTLDKINADPVLLPNITLGSEIRD<br>SCWHSSVALEQSIEFIRDSLISIRDEKDGLNRCLPDGQTLPPGRTKKPIAGVIGPGSSSVAIQVQNLLQLFDIPQIAYSATSIDLSDKTLYKYFLRVVPSDTLQARAMLDIVKRYNWTY<br>VSAVHTEGNYGESGMDAFKELAAQEGLCIAHSDKIYSNAGEKSFDRLLRKLRERLPKARVVVCFCEGMTVRGLLSAMRRLGVVGEFSLIGSDGWADRDEVIEGYEVEANGGITIK<br>LQSPEVRSFDDYFLKLRLDTNTRNPWFPEFWQHRFQCRLPGHLLENPNFKKVCTGNESLEENYVQDSKMGFVINAIYAMAHGLQNMHHALCPGHVGLCDAMKPIDGRKLLDFLI<br>KSSFVGVSGEEVWFDEKGDAPGRYDIMNLQYTEANRYDYVHVGTWHEGVLNIDDYKIQMNKSGMVRSVCSEPCLKGQIKVIRKGEVSCCWICTACKENEFVQDEFTCRACDLG<br>WWPNAELTGCEPIPVRYLEWSDIESIIAIAFSCLGILVTLFVTLIFVLYRDTPVVKSSSRELCYIILAGIFLGYVCPFTLIAKPTTTSCYLQRLLVGLSSAMCYSALVTKTNRIARILAGSKK<br>KICTRKPRFMSAWAQVIIASILISVQLTLVVTLIIMEPPMPILSYPSIKEVYLICNTSNLGVVAPVGYNGLLIMSCTYYAFKTRNVPANFNEAKYIAFTMYTTCIIWLAFVPIYFGSNYKIITT<br>CFAVSLSVTVALGCMFTPKMYIIIAKPERNVRSAFTTSDVVRMHVGDGKLPCRSNTFLNIFRRKKPGAGNANSNGKSVSWSEPGGRQAPKGQHVWQRLSVHVKTNETACNQTAV<br>IKPLTKSYQGSGKSLTFSDASTKTLYNVEEEDNTPSAHFSPPSSPSMVVHRRGPPVATTPPLPPHLTAEETPLFLADSVIPKGLPPPLPQQQPQQPPPQQPPQQPKSLMDQLQGV<br>VTNFGSGIPDFHAVLAGPGTPGNSLRSLYPPPPPPQHLQMLPLHLSTFQEESISPPGEDIDDDSERFKLLQEFVYEREGNTEEDELEEEEDLPTASKLTPEDSPALTPPSPFRDSV<br>ASGSSVPSSPVSESVLCTPPNVTYASVILRDYKQSSSTL |
| ALFA-rmGlu$_1$ - HiBiT | MVLLLILSVLLLKEDVRGSAQSTRPSRLEEELRRRLTEPDKLASSQRSVARMDGDVIIGALFSVHHQPPAEKVPERKCGEIREQYGIQRVEAMFHTLDKINADPVLLPNITLGSEIRD<br>SCWHSSVALEQSIEFIRDSLISIRDEKDGLNRCLPDGQTLPPGRTKKPIAGVIGPGSSSVAIQVQNLLQLFDIPQIAYSATSIDLSDKTLYKYFLRVVPSDTLQARAMLDIVKRYNWTY<br>VSAVHTEGNYGESGMDAFKELAAQEGLCIAHSDKIYSNAGEKSFDRLLRKLRERLPKARVVVCFCEGMTVRGLLSAMRRLGVVGEFSLIGSDGWADRDEVIEGYEVEANGGITIK<br>LQSPEVRSFDDYFLKLRLDTNTRNPWFPEFWQHRFQCRLPGHLLENPNFKKVCTGNESLEENYVQDSKMGFVINAIYAMAHGLQNMHHALCPGHVGLCDAMKPIDGRKLLDFLI<br>KSSFVGVSGEEVWFDEKGDAPGRYDIMNLQYTEANRYDYVHVGTWHEGVLNIDDYKIQMNKSGMVRSVCSEPCLKGQIKVIRKGEVSCCWICTACKENEFVQDEFTCRACDLG<br>WWPNAELTGCEPIPVRYLEWSDIESIIAIAFSCLGILVTLFVTLIFVLYRDTPVVKSSSRELCYIILAGIFLGYVCPFTLIAKPTTTSCYLQRLLVGLSSAMCYSALVTKTNRIARILAGSKK<br>KICTRKPRFMSAWAQVIIASILISVQLTLVVTLIIMEPPMPILSYPSIKEVYLICNTSNLGVVAPVGYNGLLIMSCTYYAFKTRNVPANFNEAKYIAFTMYTTCIIWLAFVPIYFGSNYKIITT<br>CFAVSLSVTVALGCMFTPKMYIIIAKPERNVRSAFTTSDVVRMHVGDGKLPCRSNTFLNIFRRKKPGAGNAGGGGSVSGWRLFKKISSGGGGNSNGKSVSWSEPGGRQAPKGQ<br>HVWQRLSVHVKTNETACNQTAVIKPLTKSYQGSGKSLTFSDASTKTLYNVEEEDNTPSAHFSPPSSPSMVVHRRGPPVATTPPLPPHLTAEETPLFLADSVIPKGLPPPLPQQQP<br>QQPPPQQPPQQPKSLMDQLQGVVTNFGSGIPDFHAVLAGPGTPGNSLRSLYPPPPPPQHLQMLPLHLSTFQEESISPPGEDIDDDSERFKLLQEFVYEREGNTEEDELEEEEDLP<br>TASKLTPEDSPALTPPSPFRDSVASGSSVPSSPVSESVLCTPPNVTYASVILRDYKQSSSTL |
| ALFA-rmGlu$_1$ - LgBiT | MVLLLILSVLLLKEDVRGSAQSTRPSRLEEELRRRLTEPDKLASSQRSVARMDGDVIIGALFSVHHQPPAEKVPERKCGEIREQYGIQRVEAMFHTLDKINADPVLLPNITLGSEIRD<br>SCWHSSVALEQSIEFIRDSLISIRDEKDGLNRCLPDGQTLPPGRTKKPIAGVIGPGSSSVAIQVQNLLQLFDIPQIAYSATSIDLSDKTLYKYFLRVVPSDTLQARAMLDIVKRYNWTY<br>VSAVHTEGNYGESGMDAFKELAAQEGLCIAHSDKIYSNAGEKSFDRLLRKLRERLPKARVVVCFCEGMTVRGLLSAMRRLGVVGEFSLIGSDGWADRDEVIEGYEVEANGGITIK<br>LQSPEVRSFDDYFLKLRLDTNTRNPWFPEFWQHRFQCRLPGHLLENPNFKKVCTGNESLEENYVQDSKMGFVINAIYAMAHGLQNMHHALCPGHVGLCDAMKPIDGRKLLDFLI<br>KSSFVGVSGEEVWFDEKGDAPGRYDIMNLQYTEANRYDYVHVGTWHEGVLNIDDYKIQMNKSGMVRSVCSEPCLKGQIKVIRKGEVSCCWICTACKENEFVQDEFTCRACDLG<br>WWPNAELTGCEPIPVRYLEWSDIESIIAIAFSCLGILVTLFVTLIFVLYRDTPVVKSSSRELCYIILAGIFLGYVCPFTLIAKPTTTSCYLQRLLVGLSSAMCYSALVTKTNRIARILAGSKK<br>KICTRKPRFMSAWAQVIIASILISVQLTLVVTLIIMEPPMPILSYPSIKEVYLICNTSNLGVVAPVGYNGLLIMSCTYYAFKTRNVPANFNEAKYIAFTMYTTCIIWLAFVPIYFGSNYKIITT<br>CFAVSLSVTVALGCMFTPKMYIIIAKPERNVRSAFTTSDVVRMHVGDGKLPCRSNTFLNIFRRKKPGAGNANSNGKSVSWSEPGGRQAPKGQHVWQRLSVHVKTNETACNQTAV<br>IKPLTKSYQGSGKSLTFSDASTKTLYNVEEEDNTPSAHFSPPSSPSMVVHRRGPPVATTPPLPPHLTAEETPLFLADSVIPKGLPPPLPQQQPQQPPPQQPPQQPKSLMDQLQGV<br>VTNFGSGIPDFHAVLAGPGTPGNSLRSLYPPPPPPQHLQMLPLHLSTFQEESISPPGEDIDDDSERFKLLQEFVYEREGNTEEDELEEEEDLPTASKLTPEDSPALTPPSPFRDSV<br>ASGSSVPSSPVSESVLCTPPNVTYASVILRDYKQSSSTLGSPPARATLEVFTLEDFVGDWEQTAAYNLDQVLEQGGVSSLLQNLAVSVTPIQRIVRSGENALKIDIHVIIPYEGLSAD<br>QMAQIEEVFKVVYPVDDHHFKVILPYGTLVIDGVTPNMLNYFGRPYEGIAVFDGKKITVTGTLWNGNKIIDERLITPDGSMLFRVTINS |
| ALFA-rmGlu$_5$ WT | MVLLLILSVLLLKEDVRGSAQSTRPSRLEEELRRRLTEPDKLQSSERRVVAHMPGDIIIGALFSVHHQPTVDKVHERKCGAVREQYGIQRVEAMLHTLERINSDPTLLPNITLGCEIR<br>DSCWHSAVALEQSIEFIRDSLISSEEEEGLVRCVDGSSSFRSKKPIVGVIGPGSSSVAIQVQNLLQLFNIPQIAYSATSMDLSDKTLFKYFMRVVPSDAQQARAMVDIVKRYNWTYV<br>SAVHTEGNYGESGMEAFKDMSAKEGICIAHSYKIYSNAGEQSFDKLLKKLRSHLPKARVVACFCEGMTVRGLLMAMRRLGLAGEFLLLGSDGWADRYDVTDGYQREAVGGITIKL<br>QSPDVKWFDDYYLKLRPETNLRNPWFQEFWQHRFQCRLEGFAQENSKYNKTCNSSLTLRTHHVQDSKMGFVINAIYSMAYGLHNMQMSLCPGYAGLCDAMKPIDGRKLLDSL<br>MKTNFTGVSGDMILFDENGDSPGRYEIMNFKEMGKDYFDYINVGSWDNGELKMDDDEVWSKKNNIIRSVCSEPCEKGQIKVIRKGEVSCCWTCTPCKENEYVFDEYTCKACQLG<br>SWPTDDLTGCDLIPVQYLRWGDPEPIAAVVFACLGLLATLFVTVIFIIYRDTPVVKSSSRELCYIILAGICLGYLCTFCLIAKPKQIYCYLQRIGIGLSPAMSYSALVTKTNRIARILAGSKK<br>KICTKKPRFMSACAQLVIAFILICIQLGIIVALFIMEPPDIMHDYPSIREVYLICNTTNLGVVTPLGYNGLLILSCTFYAFKTRNVPANFNEAKYIAFTMYTTCIIWLAFVPIYFGSNYKIITMC<br>FSVSLSATVALGCMFVPKVYIILAKPERNVRSAFTTSTVVRMHVGDGKSSSAASRSSSLVNLWKRRGSSGETLSSNGKSVTWAQNEKSTRGQHLWQRLSVHINKKENPNQTAVIK<br>PFPKSTENRGPGAAAGGGSGPGVAGAGNAGCTATGGPEPPDAGPKALYDVAEAEESFPAAARPRSPSPISTLSHLAGSAGRTDDDAPSLHSETAARSSSSQGSLMEQISSVVT<br>RFTANISELNSMMLSTAATPGPPGTPICSSYLIPKEIQLPTTMTTFAEIQPLPAIEVTGGAQGATGVSPAQETPTGAESAPGKPDLEELVALTPPSPFRDSVDSGSTTPNSPVSESAL<br>CIPSSPKYDTLIIRDYTQSSSSL |
| ALFA-rmGlu$_5$ - HiBiT | MVLLLILSVLLLKEDVRGSAQSTRPSRLEEELRRRLTEPDKLQSSERRVVAHMPGDIIIGALFSVHHQPTVDKVHERKCGAVREQYGIQRVEAMLHTLERINSDPTLLPNITLGCEIR<br>DSCWHSAVALEQSIEFIRDSLISSEEEEGLVRCVDGSSSFRSKKPIVGVIGPGSSSVAIQVQNLLQLFNIPQIAYSATSMDLSDKTLFKYFMRVVPSDAQQARAMVDIVKRYNWTYV<br>SAVHTEGNYGESGMEAFKDMSAKEGICIAHSYKIYSNAGEQSFDKLLKKLRSHLPKARVVACFCEGMTVRGLLMAMRRLGLAGEFLLLGSDGWADRYDVTDGYQREAVGGITIKL<br>QSPDVKWFDDYYLKLRPETNLRNPWFQEFWQHRFQCRLEGFAQENSKYNKTCNSSLTLRTHHVQDSKMGFVINAIYSMAYGLHNMQMSLCPGYAGLCDAMKPIDGRKLLDSL<br>MKTNFTGVSGDMILFDENGDSPGRYEIMNFKEMGKDYFDYINVGSWDNGELKMDDDEVWSKKNNIIRSVCSEPCEKGQIKVIRKGEVSCCWTCTPCKENEYVFDEYTCKACQLG<br>SWPTDDLTGCDLIPVQYLRWGDPEPIAAVVFACLGLLATLFVTVIFIIYRDTPVVKSSSRELCYIILAGICLGYLCTFCLIAKPKQIYCYLQRIGIGLSPAMSYSALVTKTNRIARILAGSKK<br>KICTKKPRFMSACAQLVIAFILICIQLGIIVALFIMEPPDIMHDYPSIREVYLICNTTNLGVVTPLGYNGLLILSCTFYAFKTRNVPANFNEAKYIAFTMYTTCIIWLAFVPIYFGSNYKIITMC<br>FSVSLSATVALGCMFVPKVYIILAKPERNVRSAFTTSTVVRMHVGDGKSSSAASRSSSLVNLWKRRGSSGGGGGSVSGWRLFKKISSGGGGETLSSNGKSVTWAQNEKSTRGQ<br>HLWQRLSVHINKKENPNQTAVIKPFPKSTENRGPGAAAGGGSGPGVAGAGNAGCTATGGPEPPDAGPKALYDVAEAEESFPAAARPRSPSPISTLSHLAGSAGRTDDDAPSLHS<br>ETAARSSSSQGSLMEQISSVVTRFTANISELNSMMLSTAATPGPPGTPICSSYLIPKEIQLPTTMTTFAEIQPLPAIEVTGGAQGATGVSPAQETPTGAESAPGKPDLEELVALTPPS<br>PFRDSVDSGSTTPNSPVSESALCIPSSPKYDTLIIRDYTQSSSSL |
| ALFA-rmGlu$_5$ - LgBiT | MVLLLILSVLLLKEDVRGSAQSTRPSRLEEELRRRLTEPDKLQSSERRVVAHMPGDIIIGALFSVHHQPTVDKVHERKCGAVREQYGIQRVEAMLHTLERINSDPTLLPNITLGCEIR<br>DSCWHSAVALEQSIEFIRDSLISSEEEEGLVRCVDGSSSFRSKKPIVGVIGPGSSSVAIQVQNLLQLFNIPQIAYSATSMDLSDKTLFKYFMRVVPSDAQQARAMVDIVKRYNWTYV<br>SAVHTEGNYGESGMEAFKDMSAKEGICIAHSYKIYSNAGEQSFDKLLKKLRSHLPKARVVACFCEGMTVRGLLMAMRRLGLAGEFLLLGSDGWADRYDVTDGYQREAVGGITIKL<br>QSPDVKWFDDYYLKLRPETNLRNPWFQEFWQHRFQCRLEGFAQENSKYNKTCNSSLTLRTHHVQDSKMGFVINAIYSMAYGLHNMQMSLCPGYAGLCDAMKPIDGRKLLDSL<br>MKTNFTGVSGDMILFDENGDSPGRYEIMNFKEMGKDYFDYINVGSWDNGELKMDDDEVWSKKNNIIRSVCSEPCEKGQIKVIRKGEVSCCWTCTPCKENEYVFDEYTCKACQLG<br>SWPTDDLTGCDLIPVQYLRWGDPEPIAAVVFACLGLLATLFVTVIFIIYRDTPVVKSSSRELCYIILAGICLGYLCTFCLIAKPKQIYCYLQRIGIGLSPAMSYSALVTKTNRIARILAGSKK<br>KICTKKPRFMSACAQLVIAFILICIQLGIIVALFIMEPPDIMHDYPSIREVYLICNTTNLGVVTPLGYNGLLILSCTFYAFKTRNVPANFNEAKYIAFTMYTTCIIWLAFVPIYFGSNYKIITMC<br>FSVSLSATVALGCMFVPKVYIILAKPERNVRSAFTTSTVVRMHVGDGKSSSAASRSSSLVNLWKRRGSSGETLSSNGKSVTWAQNEKSTRGQHLWQRLSVHINKKENPNQTAVIK<br>PFPKSTENRGPGAAAGGGSGPGVAGAGNAGCTATGGPEPPDAGPKALYDVAEAEESFPAAARPRSPSPISTLSHLAGSAGRTDDDAPSLHSETAARSSSSQGSLMEQISSVVT<br>RFTANISELNSMMLSTAATPGPPGTPICSSYLIPKEIQLPTTMTTFAEIQPLPAIEVTGGAQGATGVSPAQETPTGAESAPGKPDLEELVALTPPSPFRDSVDSGSTTPNSPVSESAL<br>CIPSSPKYDTLIIRDYTQSSSSLGSPPARATLEVFTLEDFVGDWEQTAAYNLDQVLEQGGVSSLLQNLAVSVTPIQRIVRSGENALKIDIHVIIPYEGLSADQMAQIEEVFKVVYPVDD<br>HHFKVILPYGTLVIDGVTPNMLNYFGRPYEGIAVFDGKKITVTGTLWNGNKIIDERLITPDGSMLFRVTINS |
| Chimeras ALFA-rmGlu$_5^{1ECD}$ - HiBiT or ALFA-rmGlu$_1^{5TM}$ -HiBiT | MVLLLILSVLLLKEDVRGSAQSTRPSRLEEELRRRLTEPDKLASSQRSVARMDGDVIIGALFSVHHQPPAEKVPERKCGEIREQYGIQRVEAMFHTLDKINADPVLLPNITLGSEIRD<br>SCWHSSVALEQSIEFIRDSLISIRDEKDGLNRCLPDGQTLPPGRTKKPIAGVIGPGSSSVAIQVQNLLQLFDIPQIAYSATSIDLSDKTLYKYFLRVVPSDTLQARAMLDIVKRYNWTY<br>VSAVHTEGNYGESGMDAFKELAAQEGLCIAHSDKIYSNAGEKSFDRLLRKLRERLPKARVVVCFCEGMTVRGLLSAMRRLGVVGEFSLIGSDGWADRDEVIEGYEVEANGGITIK<br>LQSPEVRSFDDYFLKLRLDTNTRNPWFPEFWQHRFQCRLPGHLLENPNFKKVCTGNESLEENYVQDSKMGFVINAIYAMAHGLQNMHHALCPGHVGLCDAMKPIDGRKLLDFLI<br>KSSFVGVSGEEVWFDEKGDAPGRYDIMNLQYTEANRYDYVHVGTWHEGVLNIDDYKIQMNKSGMVRSVCSEPCEKGQIKVIRKGEVSCCWTCTPCKENEYVFDEYTCKACQLG<br>SWPTDDLTGCDLIPVQYLRWGDPEPIAAVVFACLGLLATLFVTVIFIIYRDTPVVKSSSRELCYIILAGICLGYLCTFCLIAKPKQIYCYLQRIGIGLSPAMSYSALVTKTNRIARILAGSKK<br>KICTKKPRFMSACAQLVIAFILICIQLGIIVALFIMEPPDIMHDYPSIREVYLICNTTNLGVVTPLGYNGLLILSCTFYAFKTRNVPANFNEAKYIAFTMYTTCIIWLAFVPIYFGSNYKIITMC<br>FSVSLSATVALGCMFVPKVYIILAKPERNVRSAFTTSTVVRMHVGDGKSSSAASRSSSLVNLWKRRGSSGGGGGSVSGWRLFKKISSGGGGETLSSNGKSVTWAQNEKSTRGQ<br>HLWQRLSVHINKKENPNQTAVIKPFPKSTENRGPGAAAGGGSGPGVAGAGNAGCTATGGPEPPDAGPKALYDVAEAEESFPAAARPRSPSPISTLSHLAGSAGRTDDDAPSLHS<br>ETAARSSSSQGSLMEQISSVVTRFTANISELNSMMLSTAATPGPPGTPICSSYLIPKEIQLPTTMTTFAEIQPLPAIEVTGGAQGATGVSPAQETPTGAESAPGKPDLEELVALTPPS<br>PFRDSVDSGSTTPNSPVSESALCIPSSPKYDTLIIRDYTQSSSSL |

| Construct | Primary Structure |
|---|---|
| Chimeras ALFA-rmGlu$_1^{5ECD}$ -LgBiT or ALFA-rmGlu$_5^{1TM}$ -LgBiT | MVLLLILSVLLLKEDVRGSAQSTRPSRLEEELRRRLTEPDKLQSSERRVVAHMPGDIIIGALFSVHHQPTVDKVHERKCGAVREQYGIQRVEAMLHTLERINSDPTLLPNITLGCEIR<br>DSCWHSAVALEQSIEFIRDSLISSEEEEGLVRCVDGSSSFRSKKPIVGVIGPGSSSVAIQVQNLLQLFNIPQIAYSATSMDLSDKTLFKYFMRVVPSDAQQARAMVDIVKRYNWTYV<br>SAVHTEGNYGESGMEAFKDMSAKEGICIAHSYKIYSNAGEQSFDKLLKKLRSHLPKARVVACFCEGMTVRGLLMAMRRLGLAGEFLLLGSDGWADRYDVTDGYQREAVGGITIKL<br>QSPDVKWFDDYYLKLRPETNLRNPWFQEFWQHRFQCRLEGFAQENSKYNKTCNSSLTLRTHHVQDSKMGFVINAIYSMAYGLHNMQMSLCPGYAGLCDAMKPIDGRKLLDSL<br>MKTNFTGVSGDMILFDENGDSPGRYEIMNFKEMGKDYFDYINVGSWDNGELKMDDDEVWSKKNNIIRSVCSEPCLKGQIKVIRKGEVSCCWICTACKENEFVQDEFTCRACDLG<br>WWPNAELTGCEPIPVRYLEWSDIESIIAIAFSCLGILVTLFVTLIFVLYRDTPVVKSSSRELCYIILAGIFLGYVCPFTLIAKPTTTSCYLQRLLVGLSSAMCYSALVTKTNRIARILAGSKK<br>KICTRKPRFMSAWAQVIIASILISVQLTLVVTLIIMEPPMPILSYPSIKEVYLICNTSNLGVVAPVGYNGLLIMSCTYYAFKTRNVPANFNEAKYIAFTMYTTCIIWLAFVPIYFGSNYKIITT<br>CFAVSLSVTVALGCMFTPKMYIIIAKPERNVRSAFTTSDVVRMHVGDGKLPCRSNTFLNIFRRKKPGAGNANSNGKSVSWSEPGGRQAPKGQHVWQRLSVHVKTNETACNQTAV<br>IKPLTKSYQGSGKSLTFSDASTKTLYNVEEEDNTPSAHFSPPSSPSMVVHRRGPPVATTPPLPPHLTAEETPLFLADSVIPKGLPPPLPQQQPQQPPPQQPPQQPKSLMDQLQGV<br>VTNFGSGIPDFHAVLAGPGTPGNSLRSLYPPPPPPQHLQMLPLHLSTFQEESISPPGEDIDDDSERFKLLQEFVYEREGNTEEDELEEEEDLPTASKLTPEDSPALTPPSPFRDSV<br>ASGSSVPSSPVSESVLCTPPNVTYASVILRDYKQSSSTLGSPPARATLEVFTLEDFVGDWEQTAAYNLDQVLEQGGVSSLLQNLAVSVTPIQRIVRSGENALKIDIHVIIPYEGLSAD<br>QMAQIEEVFKVVYPVDDHHFKVILPYGTLVIDGVTPNMLNYFGRPYEGIAVFDGKKITVTGTLWNGNKIIDERLITPDGSMLFRVTINS |
| HA-rmGlu$_1$-GB1 | MVRLLLIFFPMIFLEMSILPRYPYDVPDYAMPDRKVLLAGASSQRSVARMDGDVIIGALFSVHHQPPAEKVPERKCGEIREQYGIQRVEAMFHTLDKINADPVLLPNITLGSEIRDSC<br>WHSSVALEQSIEFIRDSLISIRDEKDGLNRCLPDGQTLPPGRTKKPIAGVIGPGSSSVAIQVQNLLQLFDIPQIAYSATSIDLSDKTLYKYFLRVVPSDTLQARAMLDIVKRYNWTYVSA<br>VHTEGNYGESGMDAFKELAAQEGLCIAHSDKIYSNAGEKSFDRLLRKLRERLPKARVVVCFCEGMTVRGLLSAMRRLGVVGEFSLIGSDGWADRDEVIEGYEVEANGGITIKLQS<br>PEVRSFDDYFLKLRLDTNTRNPWFPEFWQHRFQCRLPGHLLENPNFKKVCTGNESLEENYVQDSKMGFVINAIYAMAHGLQNMHHALCPGHVGLCDAMKPIDGRKLLDFLIKSS<br>FVGVSGEEVWFDEKGDAPGRYDIMNLQYTEANRYDYVHVGTWHEGVLNIDDYKIQMNKSGMVRSVCSEPCLKGQIKVIRKGEVSCCWICTACKENEFVQDEFTCRACDLGWWP<br>NAELTGCEPIPVRYLEWSDIESIIAIAFSCLGILVTLFVTLIFVLYRDTPVVKSSSRELCYIILAGIFLGYVCPFTLIAKPTTTSCYLQRLLVGLSSAMCYSALVTKTNRIARILAGSKKKICT<br>RKPRFMSAWAQVIIASILISVQLTLVVTLIIMEPPMPILSYPSIKEVYLICNTSNLGVVAPVGYNGLLIMSCTYYAFKTRNVPANFNEAKYIAFTMYTTCIIWLAFVPIYFGSNYKIITTCFAV<br>SLSVTVALGCMFTPKMYIIIAKPERNVRSAFTTSDVVRMHAAATGSSTNNNEEEKSRLLEKENRELEKIIAEKEERVSELRHQLQSRQQLKKTN* |
| HA-rmGlu$_1$-GB2 | MVRLLLIFFPMIFLEMSILPRYPYDVPDYAMPDRKVLLAGASSQRSVARMDGDVIIGALFSVHHQPPAEKVPERKCGEIREQYGIQRVEAMFHTLDKINADPVLLPNITLGSEIRDSC<br>WHSSVALEQSIEFIRDSLISIRDEKDGLNRCLPDGQTLPPGRTKKPIAGVIGPGSSSVAIQVQNLLQLFDIPQIAYSATSIDLSDKTLYKYFLRVVPSDTLQARAMLDIVKRYNWTYVSA<br>VHTEGNYGESGMDAFKELAAQEGLCIAHSDKIYSNAGEKSFDRLLRKLRERLPKARVVVCFCEGMTVRGLLSAMRRLGVVGEFSLIGSDGWADRDEVIEGYEVEANGGITIKLQS<br>PEVRSFDDYFLKLRLDTNTRNPWFPEFWQHRFQCRLPGHLLENPNFKKVCTGNESLEENYVQDSKMGFVINAIYAMAHGLQNMHHALCPGHVGLCDAMKPIDGRKLLDFLIKSS<br>FVGVSGEEVWFDEKGDAPGRYDIMNLQYTEANRYDYVHVGTWHEGVLNIDDYKIQMNKSGMVRSVCSEPCLKGQIKVIRKGEVSCCWICTACKENEFVQDEFTCRACDLGWWP<br>NAELTGCEPIPVRYLEWSDIESIIAIAFSCLGILVTLFVTLIFVLYRDTPVVKSSSRELCYIILAGIFLGYVCPFTLIAKPTTTSCYLQRLLVGLSSAMCYSALVTKTNRIARILAGSKKKICT<br>RKPRFMSAWAQVIIASILISVQLTLVVTLIIMEPPMPILSYPSIKEVYLICNTSNLGVVAPVGYNGLLIMSCTYYAFKTRNVPANFNEAKYIAFTMYTTCIIWLAFVPIYFGSNYKIITTCFAV<br>SLSVTVALGCMFTPKMYIIIAKPERNVRSAFTTSDVVRMHAAATSTSVTSVNQASTSRLEGLQSENHRLRMKITELDKDLEEVTMQLQDTPEKKTN* |
| FLAG-rmGlu$_5$-GB1 | MVLLLILSVLLLKEDVRGSAQSDYKDDDDKSERRVVAHMPGDIIIGALFSVHHQPTVDKVHERKCGAVREQYGIQRVEAMLHTLERINSDPTLLPNITLGCEIRDSCWHSAVALEQSI<br>EFIRDSLISSEEEEGLVRCVDGSSSFRSKKPIVGVIGPGSSSVAIQVQNLLQLFNIPQIAYSATSMDLSDKTLFKYFMRVVPSDAQQARAMVDIVKRYNWTYVSAVHTEGNYGESGM<br>EAFKDMSAKEGICIAHSYKIYSNAGEQSFDKLLKKLRSHLPKARVVACFCEGMTVRGLLMAMRRLGLAGEFLLLGSDGWADRYDVTDGYQREAVGGITIKLQSPDVKWFDDYYLK<br>LRPETNLRNPWFQEFWQHRFQCRLEGFAQENSKYNKTCNSSLTLRTHHVQDSKMGFVINAIYSMAYGLHNMQMSLCPGYAGLCDAMKPIDGRKLLDSLMKTNFTGVSGDMILF<br>DENGDSPGRYEIMNFKEMGKDYFDYINVGSWDNGELKMDDDEVWSKKNNIIRSVCSEPCEKGQIKVIRKGEVSCCWTCTPCKENEYVFDEYTCKACQLGSWPTDDLTGCDLIPV<br>QYLRWGDPEPIAAVVFACLGLLATLFVTVIFIIYRDTPVVKSSSRELCYIILAGICLGYLCTFCLIAKPKQIYCYLQRIGIGLSPAMSYSALVTKTNRIARILAGSKKKICTKKPRFMSACAQ<br>LVIAFILICIQLGIIVALFIMEPPDIMHDYPSIREVYLICNTTNLGVVTPLGYNGLLILSCTFYAFKTRNVPANFNEAKYIAFTMYTTCIIWLAFVPIYFGSNYKIITMCFSVSLSATVALGCMF<br>VPKVYIILAKPERNVRSAFTTSTVVRMHAAATGSSTNNNEEEKSRLLEKENRELEKIIAEKEERVSELRHQLQSRQQLKKTN* |
| FLAG-rmGlu$_5$-GB2 | MVLLLILSVLLLKEDVRGSAQSDYKDDDDKSERRVVAHMPGDIIIGALFSVHHQPTVDKVHERKCGAVREQYGIQRVEAMLHTLERINSDPTLLPNITLGCEIRDSCWHSAVALEQSI<br>EFIRDSLISSEEEEGLVRCVDGSSSFRSKKPIVGVIGPGSSSVAIQVQNLLQLFNIPQIAYSATSMDLSDKTLFKYFMRVVPSDAQQARAMVDIVKRYNWTYVSAVHTEGNYGESGM<br>EAFKDMSAKEGICIAHSYKIYSNAGEQSFDKLLKKLRSHLPKARVVACFCEGMTVRGLLMAMRRLGLAGEFLLLGSDGWADRYDVTDGYQREAVGGITIKLQSPDVKWFDDYYLK<br>LRPETNLRNPWFQEFWQHRFQCRLEGFAQENSKYNKTCNSSLTLRTHHVQDSKMGFVINAIYSMAYGLHNMQMSLCPGYAGLCDAMKPIDGRKLLDSLMKTNFTGVSGDMILF<br>DENGDSPGRYEIMNFKEMGKDYFDYINVGSWDNGELKMDDDEVWSKKNNIIRSVCSEPCEKGQIKVIRKGEVSCCWTCTPCKENEYVFDEYTCKACQLGSWPTDDLTGCDLIPV<br>QYLRWGDPEPIAAVVFACLGLLATLFVTVIFIIYRDTPVVKSSSRELCYIILAGICLGYLCTFCLIAKPKQIYCYLQRIGIGLSPAMSYSALVTKTNRIARILAGSKKKICTKKPRFMSACAQ<br>LVIAFILICIQLGIIVALFIMEPPDIMHDYPSIREVYLICNTTNLGVVTPLGYNGLLILSCTFYAFKTRNVPANFNEAKYIAFTMYTTCIIWLAFVPIYFGSNYKIITMCFSVSLSATVALGCMF<br>VPKVYIILAKPERNVRSAFTTSTVVRMHAAATSTSVTSVNQASTSRLEGLQSENHRLRMKITELDKDLEEVTMQLQDTPEKKTN* |

**Table S1. Primary Structure of Constructs Used in Assays.**
Mutations differ between CODA-RET and GABA$_B$-tail assays in the following ways: for mGlu$_1$ glutamate-binding deficient mutants T188A and R78L, respectively, for mGlu$_5$ glutamate-binding deficient mutants, T174A and R68E, respectively, for mGlu$_1$ G$\alpha_q$-coupling deficient mutants, F781D and F781S, respectively and for mGlu$_5$ G$\alpha_q$-coupling deficient mutants, F767D and F767S, respectively.

| Fig Ref | Construct | EC50 µM (±SEM) | Emax AU (±SEM) | n | Fig Ref | Construct | EC50 µM (±SEM) | Emax AU (±SEM) | n |
|---|---|---|---|---|---|---|---|---|---|
| 1c | $mGlu_{1/1}$ WT CR | 13.91 (13.34-14.48) | 56.99 (55.63-58.35) | 3 | 1c | $mGlu_{1/5}$ WT CR | 8.42 (6.82-10.02) | 56.51 (53.20-59.82) | 3 |
| 1c | $mGlu_{5/5}$ WT CR | 8.81 (7.21-10.41) | 50.10 (48.71-51.49) | 3 | 1g | $mGlu_{1/1}$ WT GT | 12.9 (12.3-13.5) | 73.6 (70.8-76.4) | 4 |
| 1g | $mGlu_{1/5}$ WT GT | 1.51 (1.35-1.70) | 96.8 (90.8-102.8) | 3 | 1g | $mGlu_{5/5}$ WT GT | 0.96 (0.91-1.00) | 83.1 (78.0-88.2) | 4 |
| 2a | $mGlu_{1/1}$ WT | 10.94 (8.33-13.55) | 60.74 (57.29-64.19) | 3 | 2a | $mGlu_{1/1}$ Trans | N/A | 4.76 (1.77-7.74) | 3 |
| 2b | $mGlu_{5/5}$ WT | 11.24 (9.12-13.36) | 53.21 (49.44-56.98) | 4 | 2b | $mGlu_{5/5}$ Trans | 3198 (3009-3387) | 24.29 (22.93-25.66) | 4 |
| 2c | $mGlu_{1/5}$ WT | 6.53 (5.48-7.58) | 53.97 (50.41-57.53) | 3 | 2c | Trans 5=>1 | 3256 (2431-4081) | 20.79 (17.97-23.60) | 3 |
| 2c | Trans 1=>5 | 5128 (2943-7313) | 18.33 (15.61-21.05) | 3 | 3a | mGlu1/1 WT | 13.90 (9.86-17.94) | 56.99 (55.63-58.34) | 3 |
| 3a | $mGlu_{1/1}$ Cis | N/A | 6.97 (5.65-8.30) | 3 | 3b | $mGlu_{5/5}$ WT | 8.80 (7.19-10.40) | 50.11 (48.72-51.50) | 3 |
| 3b | $mGlu_{5/5}$ Cis | 3181 (1574-4787) | 27.56 (26.86-28.26) | 3 | 3c | $mGlu_{1/5}$ WT | 10.52 (8.34-12.71) | 57.03 (54.03-60.06) | 10 |
| 3c | Cis 1 | 1094 (979-1208) | 38.96 (36.64-41.28) | 10 | 3c | Cis 5 | 124.3 (77.8-170.9) | 10.84 (10.14-11.54) | 9 |
| 4a | $mGlu_{1/1}$ WT | 26.39 (12.87-39.91) | 63.15 (56.07-70.23) | 7 | 4a | 1 $G\alpha_q$ only | 32.26 (21.06-43.46) | 45.15 (38.57-51.72) | 7 |
| 4b | $mGlu_{5/5}$ WT | 15.35 (10.20-20.50) | 49.91 (44.52-55.30) | 7 | 4b | 5 $G\alpha_q$ only | 16.75 (10.71-22.79) | 42.93 (38.64-47.22) | 7 |
| 4c | $mGlu_{1/5}$ WT CR | 8.95 (5.43-12.47) | 56.64 (47.72-65.56) | 12 | 4c | $G\alpha_q$ 1 only CR | 10.42 (6.81-14.03) | 57.35 (48.96-65.74) | 12 |
| 4c | $G\alpha_q$ 5 only CR | 19.13 (12.42-25.84) | 32.69 (26.42-38.95) | 11 | 4e | $mGlu_{1/5}$ WT GT | 1.29 (1.15-1.45) | 99.3 (86.9-111.7) | 3 |
| 4e | $G\alpha_q$ 1 only GT | 2.14 (1.91-2.40) | 29.4 (21.3-37.5) | 3 | 4e | $G\alpha_q$ 5 only GT | 2.29 (1.91-2.75) | 73.6 (66.4-80.8) | 3 |
| 5a | $mGlu_{1/1}$ WT | 32.14(19.76-44.52) | 69.7 (63.44-75.96) | 3 | 5a | $mGlu_5^{1ECD}/mGlu_5^{WT}$ | 15.54 (13.40-17.68) | 46.89 (44.53-49.25) | 3 |
| 5a | $mGlu_5^{1ECD}$ $G\alpha_q$ only | 18.32 (10.44-26.21) | 36.68 (33.27-40.09) | 3 | 5a | 5 WT $G\alpha_q$ Only | 15.65(10.14-21.16) | 34.20 (31.16-37.24) | 3 |
| 5b | $mGlu_{1/1}$ WT | 32.60 (20.74-44.46) | 77.83(74.18-81.48) | 3 | 5b | $mGlu_1^{5ECD}/mGlu_1^{WT}$ | 25.49 (17.03-33.95) | 73.22(63.36-83.09) | 3 |
| 5b | $mGlu_1^{5ECD}$ $G\alpha_q$ only | 26.23 (11.76-40.70) | 56.13 (46.36-65.90) | 3 | 5b | 1 WT $G\alpha_q$ Only | 31.88 (14.52-49.24) | 54.57 (49.81-59.34) | 3 |
| 5c | $mGlu_{1/1}$ WT | 30.2 (22.58-37.82) | 73.35 (65.23-81.47) | 3 | 5c | $mGlu_5^{1TM}/mGlu_5^{WT}$ | 19.47 (12.48-26.46) | 64.89 (52.25-77.53) | 3 |
| 5c | $mGlu_5^{1TM}$ $G\alpha_q$ only | 21.37 (8.94-33.80) | 58.61 (50.21-67.02) | 3 | 5c | 5 WT $G\alpha_q$ Only | 27.84(24.81-30.87) | 39.63 (29.91-49.35) | 3 |
| 5d | $mGlu_{1/1}$ WT | 34.00 (12.88-55.12) | 70.99 (62.76-79.22) | 3 | 5d | $mGlu_1^{5TM}/mGlu_1^{WT}$ | 28.51 (2.78-54.24) | 70.87 (62.92-78.82) | 3 |
| 5d | $mGlu_1^{5TM}$ $G\alpha_q$ only | 43.20 (11.71-74.69) | 38.57 (34.86-42.28) | 3 | 5d | 1 WT $G\alpha_q$ Only | 19.02 (2.95-35.09) | 58.25 (50.02-66.48) | 3 |
| 6a | $mGlu_{1/1}$ WT +4578 | 17.79 (7.90-27.68) | 80.72 (75.22-86.22) | 3 | 6a | $mGlu_{1/1}$ WT+FITM | ND | 8.13 (6.74-9.52) | 3 |
| 6b | $mGlu_{5/5}$ WT+4578 | 12.65 (10.17-15.13) | 47.26 (45.06-49.46) | 3 | 6b | $mGlu_{5/5}$ WT+FITM | 12.65 (10.17-15.13) | 47.26 (45.06-49.46) | 3 |
| 6c | $mGlu_{1/5}$ WT +4578 | 7.81 (2.26-13.35) | 66.56 (57.77-75.35) | 9 | 6c | $mGlu_{1/5}$ WT +FITM | 23.84 (14.12-33.56) | 36.82 (30.5-43.13) | 6 |
| 6d | $G\alpha_q$ 1 only+4578 | 7.83 (3.08-12.58) | 70.52 (62.12-78.92) | 9 | 6d | $G\alpha_q$ 1 only+FITM | ND | 2.72 (-0.01-5.45) | 6 |
| 6e | $G\alpha_q$ 5 only+4578 | 14.12 (5.91-22.33) | 23.9 (20.13-27.67) | 9 | 6e | $G\alpha_q$ 5 only+FITM | 31.75 (22.85-40.64) | 41.72 (36.29-47.15) | 6 |
| 6f | $mGlu_{1/5}$ WT+MTEP | 13.72 (7.81-19.63) | 50.56 (41.41-59.71) | 5 | 6g | $G\alpha_q$ 1 only+MTEP | 12.48 (8.96-16.00) | 49.1 (43.20-55.20) | 5 |
| 6h | $G\alpha_q$ 5 only+MTEP | 32.41 (13.86-50.96) | 8.60 (5.04-12.17) | 4 | 7a | $mGlu_{1/1}$ WT | 37.09 (9.14-65.04) | 69.80 (60.98-78.63) | 5 |
| 7a | $mGlu_{1/1}$ WT + 4578 | 11.82 (5.36-18.28) | 89.31(81.19-97.44) | 4 | 7b | $mGlu_1^{Y672V}/mGlu_1^{Y672V}$ | 25.6 (4.25-46.95) | 62.07 (54.26-69.88) | 5 |
| 7b | $mGlu_1^{Y672V}/mGlu_1^{Y672V}$ + 4578 | 15.57 (4.77-26.37) | 63.55 (54.88-72.22) | 4 | 7c | $mGlu_1^{Y672V,F781D}/mGlu_1$ | 19.29 (7.51-31.07) | 42.51 (36.87-48.14) | 4 |
| 7c | $mGlu_1^{Y672V,F781D}/mGlu_1$ +4578 | 15.99 (4.58-27.40) | 68.39 (62.69-74.10) | 4 | 7d | $mGlu_1^{Y672V}/mGlu_1^{F781D}$ | 29.86 (11.72-48.00) | 41.88 (32.08-51.68) | 4 |
| 7d | $mGlu_1^{Y672V}/mGlu_1^{F781D}$ + 4578 | 27.62 (10.84-44.40) | 42.6 (33.45-51.75) | 4 | 7e | $mGlu_{1/1}$ WT | 36.05 (23.92-48.18) | 66.08 (58.26-73.90) | 4 |
| 7e | $mGlu_{1/1}$ WT + FITM | 41.51 (26.97-56.06) | 21.58 (19.51-23.65) | 4 | 7f | $mGlu_1^{G665F}/mGlu_1^{G665F}$ | 47.23 (35.10-59.36) | 37.80 (34.61-40.99) | 4 |
| 7f | $mGlu_1^{G665F}/mGlu_1^{G665F}$+ FITM | 47.32 (35.58-59.06) | 36.86 (33.89-39.83) | 4 | 7g | $mGlu_1^{G665F,F781D}/mGlu_1$ | 15.99 (4.58-27.40) | 68.39 (62.69-74.10) | 4 |
| 7g | $mGlu_1^{G665F,F781D}/mGlu_1$ +FITM | 36.84 (22.63-51.05) | 16.83 (11.26-22.40) | 4 | 7h | $mGlu_1^{G665F}/mGlu_1^{F781D}$ | 41.96 (36.03-47.89) | 22.25 (14.44-30.06) | 4 |
| 7h | $mGlu_1^{G665F}/mGlu_1^{F781D}$ +FITM | 47.36 (33.51-61.21) | 23.26 (15.52-31.00) | 4 | Ext 1a | $mGlu_1$ WT | 2.37 (1.95-2.79) | 1.37 x $10^{-6}$ (0.84-1.91) | 4 |
| Ext 1a | $mGlu_1$-HiBiT | 2.69 (2.37-3.01) | 1.35 x $10^{-6}$ (0.90-1.80) | 4 | Ext 1a | $mGlu_1$-LgBiT | 2.54 (2.00-3.08) | 1.31 (1.04-1.57) | 4 |
| Ext 1c | $mGlu_5$ WT | 1.66 (0.60-2.72) | 1.13 x $10^{-6}$ (0.46-1.81) | 3 | Ext 1c | $mGlu_5$-HiBiT | 2.97 (0.27-5.68) | 1.09 x $10^{-6}$ (0.36-1.82) | 3 |
| Ext 1c | $mGlu_5$-LgBiT | 2.24 (1.44-3.04) | 0.92 x $10^{-6}$ (0.49-1.35) | 3 | Ext 1e | $mGlu_1$ WT | 6.62 (-0.25 – 13.49) | 3.36 x $10^{-5}$ (1.47-5.24) | 4 |
| Ext 1e | $mGlu_1^{T188A}$ | ND | -2.1 x $10^{-4}$ (-4.41-0.25) | 4 | Ext 1e | $mGlu_1^{F781D}$ | ND | -0.37 x $10^{-4}$ (-4.3-3.6) | 4 |
| Ext 1e | $mGlu_1^{T188A,F781D}$ | ND | -0.18 x $10^{-4}$ (-3.7-0.05) | 4 | Ext 1e | $mGlu_1$-HiBiT | 7.42 (-0.21-15.05) | 3.66 x $10^{-5}$ (0.94-6.37) | 3 |

| Fig Ref | Construct | EC50 µM (±SEM) | Emax AU (±SEM) | n | Fig Ref | Construct | EC50 µM (±SEM ) | Emax AU (±SEM) | n |
|---|---|---|---|---|---|---|---|---|---|
| Ext 1e | $mGlu_1^{T188A}$-HiBiT | ND | $-3.2 \times 10^{-4}$ (-4.7-(-)1.6) | 3 | Ext 1e | $mGlu_1^{F781D}$-HiBiT | ND | $-3.3 \times 10^{-4}$ (-6.3- (-) 0.2) | 3 |
| Ext 1e | $mGlu_1^{T188A,F781D}$-HiBiT | ND | $-1.0 \times 10^{-4}$ (-1.9-(-)0.2) | 3 | Ext 1e | $mGlu_1$-LgBiT | 8.92 (-0.74-18.6) | $5.02 \times 10^{6}$ (2.07-7.97) | 4 |
| Ext 1e | $mGlu_1^{T188A}$-LgBiT | ND | $-1.5 \times 10^{-4}$ (-3.2 - 0.01) | 4 | Ext 1e | $mGlu_1^{F781D}$-LgBiT | ND | $-0.9 \times 10^{-4}$ (-3.4 – 1.5) | 4 |
| Ext 1e | $mGlu_1^{T188A,F781D}$-LgBiT | ND | $-2.1 \times 10^{-4}$ (-3.2- (-)1.0) | 4 | Ext 1f | $mGlu_5$ WT | 1.48 (-0.31 – 3.27) | $1.95 \times 10^{-5}$ (-0.3-4.24) | 4 |
| Ext 1f | $mGlu_5^{T174A}$ | ND | $2.1 \times 10^{-4}$ (0.4-3.7) | 3 | Ext 1f | $mGlu_5^{F767D}$ | ND | $-1.2 \times 10^{-4}$ (-3.2-0.7) | 3 |
| Ext 1f | $mGlu_5^{T174A,F767D}$ | ND | $-1.2 \times 10^{-4}$ (-2.3-(-)0.06) | 4 | Ext 1f | $mGlu_5$-HiBiT | 7.53 (-0.91-16.0) | $3.00 \times 10^{-5}$ (2.42-3.59) | 3 |
| Ext 1f | $mGlu_5^{T174A}$-HiBiT | ND | $2.8 \times 10^{-4}$ (0.3-5.2) | 3 | Ext 1f | $mGlu5_{F781D}$-HiBiT | ND | $-3.4 \times 10^{-4}$ (-8.1-1.4) | 3 |
| Ext 1f | $mGlu_5^{T174A,F767D}$-HiBiT | ND | $-2.4 \times 10^{-4}$ (-4.3-(-)0.5) | 3 | Ext 1f | $mGlu_5$-LgBiT | 3.69 (-0.21-7.59) | $4.33 \times 10^{6}$ (2.43-6.23) | 3 |
| Ext 1f | $mGlu_5^{T174A}$-LgBiT | ND | $-1.4 \times 10^{-4}$ (-4.6 - 1.9) | 3 | Ext 1f | $mGlu_5^{F767D}$-LgBiT | ND | $-4.1 \times 10^{-4}$ (-7.8 – (-)0.3) | 3 |
| Ext 1f | $mGlu_5^{T174A,F767D}$-LgBiT | ND | $-0.3 \times 10^{-4}$ (-1.2- 0.6) | 3 | Ext 2a | $mGlu_{1/5}$ WT | 10.3 (6.8-13.8) | 52.6 (45.1-60.1) | 4 |
| Ext 2a | $mGlu_{1/5}$ WT + $mGlu_1$ WT | 13.4 (9.5-17.3) | 6.59 (4.63-8.55) | 4 | Ext 2a | $mGlu_{1/5}$ WT + $mGlu_5$ WT | 34.3 (18.2-50.4) | 4.51 (3.42-5.61) | 4 |
| Ext 3a | $mGlu_1$-GB1/GB2 WT | 12.9 (12.3-13.5) | 73.6 (70.8-76.4) | 4 | Ext 3a | $mGlu_1^{R78L}$/$mGlu_1^{F781S}$-GB1/GB2 | ND | 28.0 (24.1-31.9) | 3 |
| Ext 3b | $mGlu_5$-GB1/GB2 WT | 0.96 (0.91-1.00) | 83.1 (78.0-88.2) | 4 | Ext 3b | $mGlu_5^{R68E}$/$mGlu_5^{F767S}$-GB1/GB2 | 141 (123-162) | 89.9 (83.3-96.5) | 3 |
| Ext 3c | $mGlu_{1/5}$-GB1/GB2 WT | 1.58 (1.55-1.62) | 94.5 (89.6-99.6) | 3 | Ext 3c | $mGlu1_{R78L}$/$5_{F767S}$-GB1/GB2 | 251 (234-269) | 134.9 (119.8-150.0) | 3 |
| Ext 3c | $mGlu_1^{F781S}$ /$mGlu_5^{R68E}$-GB1/GB2 | 417 (380-457) | 18.2 (16.6-19.8) | 3 | Ext 3d | $mGlu_1^{R78L,F781S}$/$mGlu_1^{WT}$-GB1/GB2 | ND | 28.9 (24.9=32.9) | 3 |
| Ext 3e | $mGlu_1^{R78L,F781}$/$mGlu_5$-GB1/GB2 | ND | ND | 3 | Ext 3e | $mGlu_1^{WT}$/$mGlu_5^{R68E,F767S}$-GB1/GB2 | 426 (371-490) | 62.1 (52.5-71.7) | 3 |
| Ext 4a | $mGlu_{1/5}$-GB1/GB2 + DMSO | 1.28 (1.15-1.45) | 99.3 (86.9-111.7) | 3 | Ext 4a | $mGlu_{1/5}$-GB1/GB2 + FITM | 2.29 (2.09-2.51) | 59.2 (49.6-68.8) | 3 |
| Ext 4a | $mGlu_{1/5}$-GB1/GB2 + 4578 | 0.78 (0.76-0.79) | 112.7 (100.5-124.9) | 3 | Ext 4b | $mGlu_1$/$mGlu_5^{F767S}$-GB1/GB2 + DMSO | 2.29 (1.91-2.75) | 73.6 (66.4-80.8) | 3 |
| Ext 4b | $mGlu_1$/$mGlu_5^{F767S}$-GB1/GB2 + FITM | ND | 0.7 (-0.1-0.8) | 3 | Ext 4b | $mGlu_1$/$mGlu_5^{F767S}$-GB1/GB2 + 4578 | 0.89 (0.79-1.00) | 108.9 (102.6- 115.2) | 3 |
| Ext 4c | $mGlu_1^{F781S}$/$mGlu_5$-GB1/GB2+ DMSO | 2.14 (1.91-2.40) | 29.4 (21.3-37.5) | 3 | Ext 4c | $mGlu_1^{F781S}$/$mGlu_5$-GB1/GB2 + FITM | 1.74 (1.70-1.78) | 105.5 (99.8-111.2) | 3 |
| Ext 4c | $mGlu_1^{F781S}$/$mGlu_5$-GB1/GB2 + 4578 | 1.41 (1.12-1.78) | 14.0 (6.6-21.4) | 3 | Ext 5a | $mGlu_{1/5}$* WT | 5.84 (5.33-6.35) | 48.51 (47.02-50.00) | 3 |
| Ext 5b | $mGlu_{1/5}$* CisTrans | 10.3 (5.9-14.6) | 57.95 (49.38-66.52) | 9 | Ext 5b | $G\alpha_q$ 1* only | 11.7 (6.8-16.6) | 56.73 (49.43-64.03) | 9 |
| Ext 5b | $G\alpha_q$ 5* only | 18.1 (11.5-24.7) | 33.21 (27.31-39.11) | 9 | Ext 5c | $G\alpha_q$ 1* only+4578 | 6.55 (4.37-8.74) | 71.15 (62.47-79.83) | 6 |
| Ext 5c | $G\alpha_q$ 1* only+FITM | NA (N/A) | 0.60 (-2.67-3.87) | 4 | Ext 5d | $G\alpha_q$ 5* only+4578 | 10.48 (6.77-14.19) | 23.71 (16.06-31.36) | 6 |
| Ext 5d | $G\alpha_q$ 5* only+FITM | 27.83 (22.76-32.90) | 42.44 (33.98-50.90) | 4 | Ext 5e | $mGlu_{1/5}$* CisTrans+MTEP | 12.64 (7.41-17.87) | 45.2 (33.31-57.09) | 5 |
| Ext 5f | $G\alpha_q$ 1* only+MTEP | 16.03 (9.87-22.19) | 52.12 (37.02-67.22) | 5 | Ext 5g | $G\alpha_q$ 5* only+MTEP | NA (NA) | 6.20 (2.70-9.70) | 5 |

**Table S2. $EC_{50}$'s and $E_{max}$'s for All Assays.**
In parentheses, ±SEM for $EC_{50}$ and $E_{max}$ and number for repeats (n). CR = CODA-RET assay; GT = $GABA_B$-tail assay.

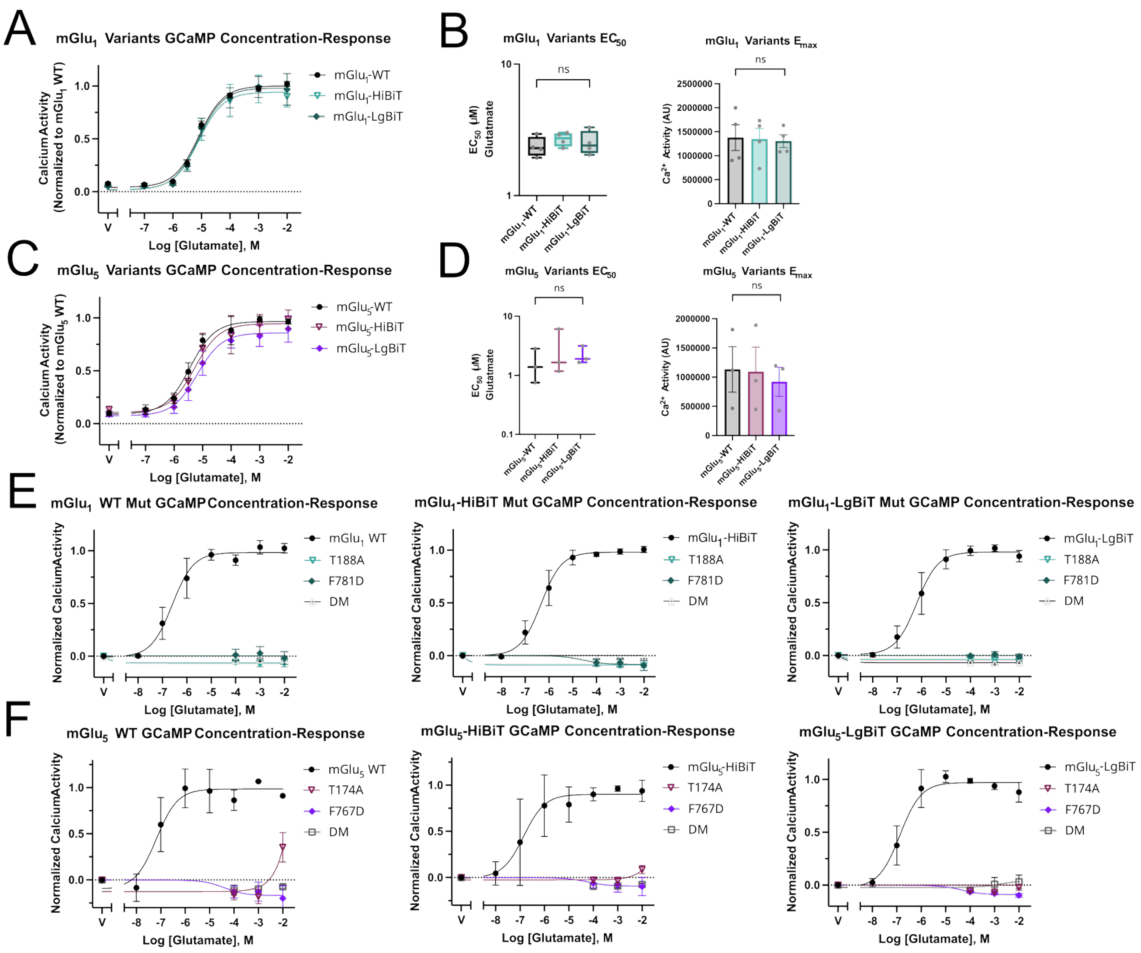

**Figure S1. Validation of Variants Using GCaMP.**

(A) Left shows a concentration-response curve of mGlu$_1$ WT with vehicle (black), mGlu$_1$-HiBiT (turquoise), and mGlu$_1$-LgBiT (teal).

(B) Left shows corresponding EC$_{50}$: ns 0.27 = and ns = 0.64. Rights shows corresponding E$_{max}$: ns = 0.93 and ns= 0.82.

(C) Left shows a concentration-response curve of mGlu$_5$ WT with vehicle (black), mGlu$_5$-HiBiT (magenta), and mGlu$_5$-LgBiT (purple).

(D) Left shows corresponding EC$_{50}$: ns 0.48 = and ns = 0.49. Rights shows corresponding E$_{max}$: ns = 0.95 and ns= 0.67.

(E) Concentration-response curves for mGlu$_1$ WT, mGlu$_1$-HiBiT, and mGlu$_1$-LgBiT (left to right-black) compared with orthosteric binding mutant T188A (turquoise), G$\alpha_q$-binding mutant F781D (teal), and double mutant (DM) T188A/F781D (grey).

(F) Concentration-response curves for mGlu$_5$ WT, mGlu$_5$-HiBiT, and mGlu$_5$-LgBiT (left to right-black) compared with orthosteric binding mutant T174A (magenta), G$\alpha_q$-binding mutant F767D (purple), and double mutant (DM) T174A/F767D (grey). Symbols represent the mean drug-induced GCAMP fluorescence and error bars represent ± SEM. The exact number of ‘n’ independent experiments and technical replicates are reported in Table S2.

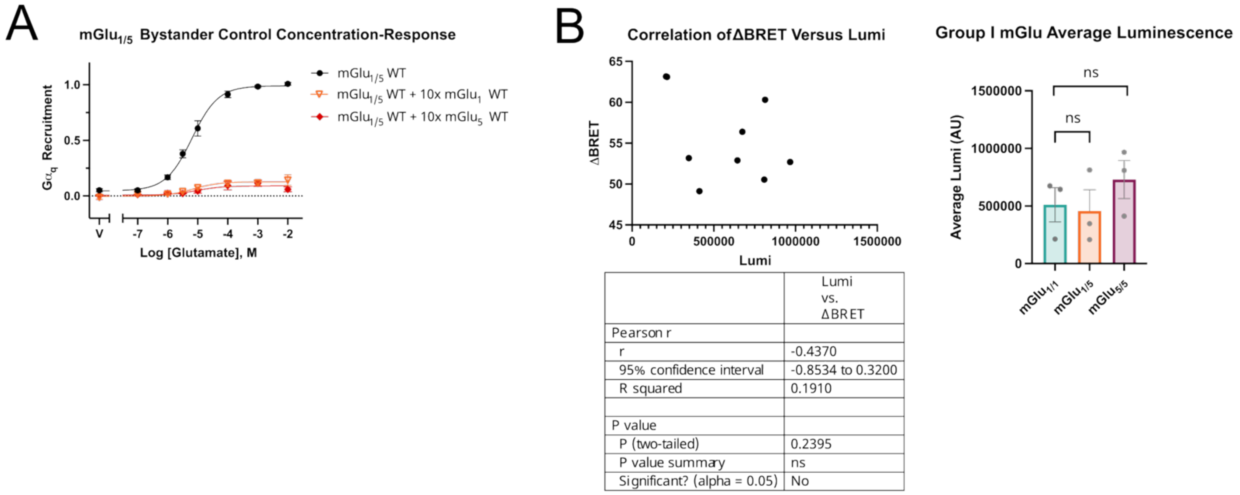

| | Lumi vs. ΔBRET |
|---|---|
| Pearson r | |
| r | -0.4370 |
| 95% confidence interval | -0.8534 to 0.3200 |
| R squared | 0.1910 |
| P value | |
| P (two-tailed) | 0.2395 |
| P value summary | ns |
| Significant? (alpha = 0.05) | No |

**Figure S2. CODA-RET Assay Validation.**
(A) Concentration-response curve of $mGlu_{1/5}$ WT (50 ng $mGlu_1$-LgBiT and 100 ng $mGlu_5$-HiBiT) (black), with 500 ng of $mGlu_1$ WT (light orange), or with 1000 ng $mGlu_5$ WT (dark orange).
(B) Left shows a correlation plot of ΔBRET versus Luminescence showing an r = -0.4370 and p = 0.24. Rights shows a plot of average luminescence for the data from Figure 1. ns = 0.83 and 0.38.

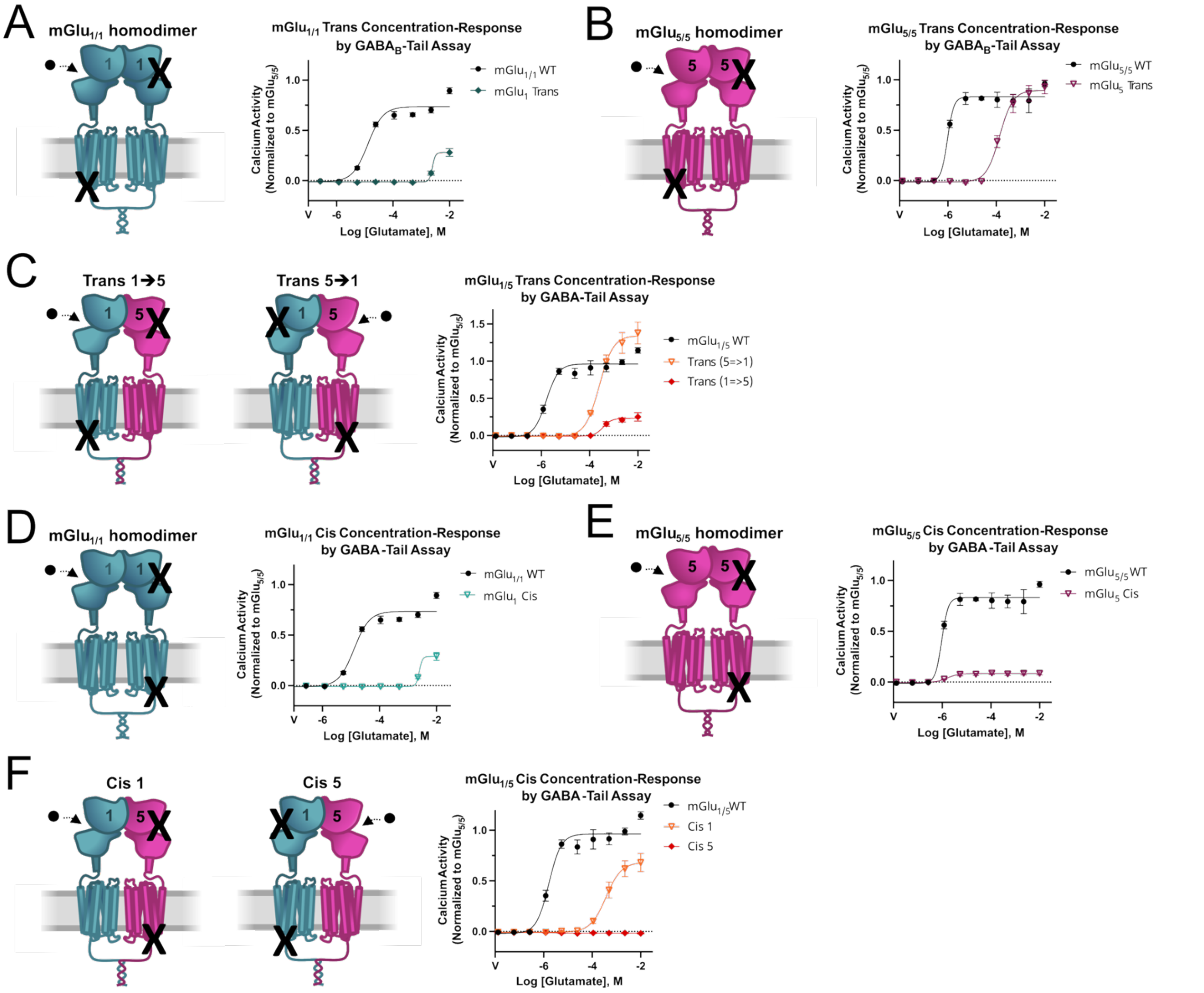

**Figure S3. $GABA_B$-tails Assay Confirmation of Trans- and Cis-activation.**

(A) Left shows schematic of R78L and F781S mutations on one $mGlu_1$-GB1 protomer paired with $mGlu_1$-GB2 WT (cyan), restricting to transactivation. Right shows a concentration-response curve of $mGlu_{1/1}$-GB1/GB2 (black) versus $mGlu_{1/1}$ in trans (cyan) by $GABA_B$-tail assay.

(B) Left shows schematic of R68E and F767S mutations on one $mGlu_5$-GB1 protomer paired with $mGlu_5$-GB2 WT (cyan), restricting to transactivation. Right shows a concentration-response curve of $mGlu_{5/5}$-GB1/GB2 WT (black) versus $mGlu_{1/1}$-GB1/GB2 in trans (cyan) by $GABA_B$-tail assay.

(C) Left shows schematic of R78L mutation on $mGlu_1$-GB1 protomer (cyan) and F767S mutation on $mGlu_5$-GB2 protomer (magenta), restricting to transactivation from $mGlu_5$ to $mGlu_1$. Middle shows schematic of F781S mutation on $mGlu_1$-GB1 protomer (cyan) and R68E mutation on $mGlu_5$-GB2 protomer (magenta), restricting to transactivation from $mGlu_1$ to $mGlu_5$. Right shows a concentration-response curve of $mGlu_{1/5}$-GB1/GB2 (black) versus $mGlu_{1/5}$-GB1/GB2 in trans from 5 to 1 (light orange) versus $mGlu_{1/5}$ in trans from 1 to 5 (dark orange) by $GABA_B$-tail assay.

(D) Left shows schematic of R78L and F781S mutations on one $mGlu_1$-GB1 protomer paired with $mGlu_1$-GB2 WT (cyan), restricting to cis activation. Right shows a concentration-response curve of $mGlu_{1/1}$-GB1/GB2 WT (black) versus $mGlu_{1/1}$ -GB1/GB2 in cis (cyan) by $GABA_B$-tail assay.

(E) Left shows schematic of R68E and F767S mutations on one mGlu$_5$-GB1 protomer paired with mGlu$_5$-GB2 (magenta), restricting to cis activation. Right shows a concentration-response curve of mGlu$_{5/5}$-GB1/GB2 (black) versus mGlu5-GB1/5-GB2 in cis (magenta) by GABA$_B$-tail assay.

(F) Left shows schematic of R68E and F767S mutations on mGlu$_5$ protomer (magenta) paired with mGlu$_1$ WT, restricting to cis activation through mGlu$_1$. Middle shows R78L and F781S mutations on mGlu-GB1 protomer (cyan) paired with mGlu$_5$-GB2 WT (magenta), restricting to cis activation through mGlu$_5$. Right shows a concentration-response curve of mGlu$_{1/5}$-GB1/GB2 (black) versus mGlu$_{1/5}$-GB1/GB2 in cis through mGlu$_1$ (light orange) versus mGlu$_{1/5}$-GB1/GB2 in cis through mGlu5 (dark orange) by GABA$_B$-tail assay. The exact number of ‘n’ independent experiments and technical replicates are reported in Table S2.

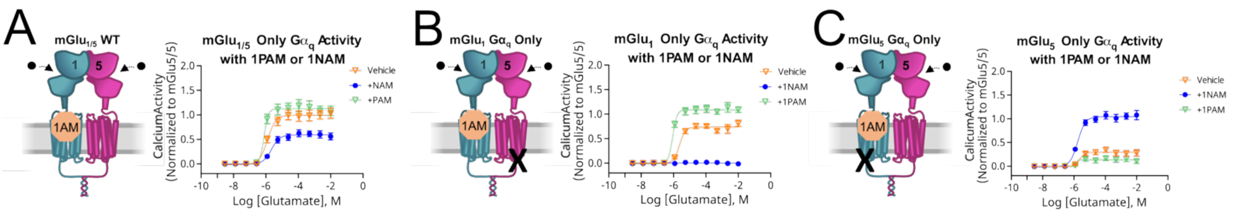

**Figure S4. GABA$_B$-tail Assay Confirmation of Inversion of mGlu$_1$ PAM and NAM Signaling.**
(A) Left shows a schematic of mGlu$_{1/5}$-GB1/GB2 with a 1PAM bound to the allosteric binding pocket of mGlu$_1$. Right shows a concentration-response curve of mGlu$_{1/5}$-GB1/GB2 with vehicle (orange), with PAM (100 nM VU6024578-green), or with NAM (100 nM FITM-blue).
(B) Left shows a schematic of mGlu$_{1/5}$-GB1/GB2 with a G$\alpha_q$-binding point mutation (F767S) in mGlu$_5$ with a 1PAM bound to the allosteric binding pocket of mGlu$_1$. Right shows a concentration-response curve of mGlu$_{1/5}$-GB1/GB2 with vehicle (grey), mGlu$_1$/mGlu5$^{F767S}$ with vehicle (orange), mGlu$_1$/mGlu5$^{F767S}$ with PAM (100 nM VU6024578-green), or mGlu$_1$/mGlu5$^{F767S}$ with NAM (100 nM FITM-blue).
(C) Left shows a schematic of mGlu$_{1/5}$-GB1/GB2 with a G$\alpha_q$-binding point mutation (F781S) in mGlu$_1$ with a 1PAM bound to the allosteric binding pocket of mGlu$_1$. Middle shows a concentration-response curve of mGlu$_{1/5}$-GB1/GB2 (grey) with vehicle, mGlu$_1^{F781S}$/mGlu$_5$ with vehicle (orange), mGlu$_1^{F781S}$/mGlu$_5$ with PAM (100 nM VU6024578-green), or mGlu$_1^{F781S}$/mGlu$_5$ with NAM (100 nM FITM-blue). The exact number of ‘n’ independent experiments and technical replicates are reported in Table S2.

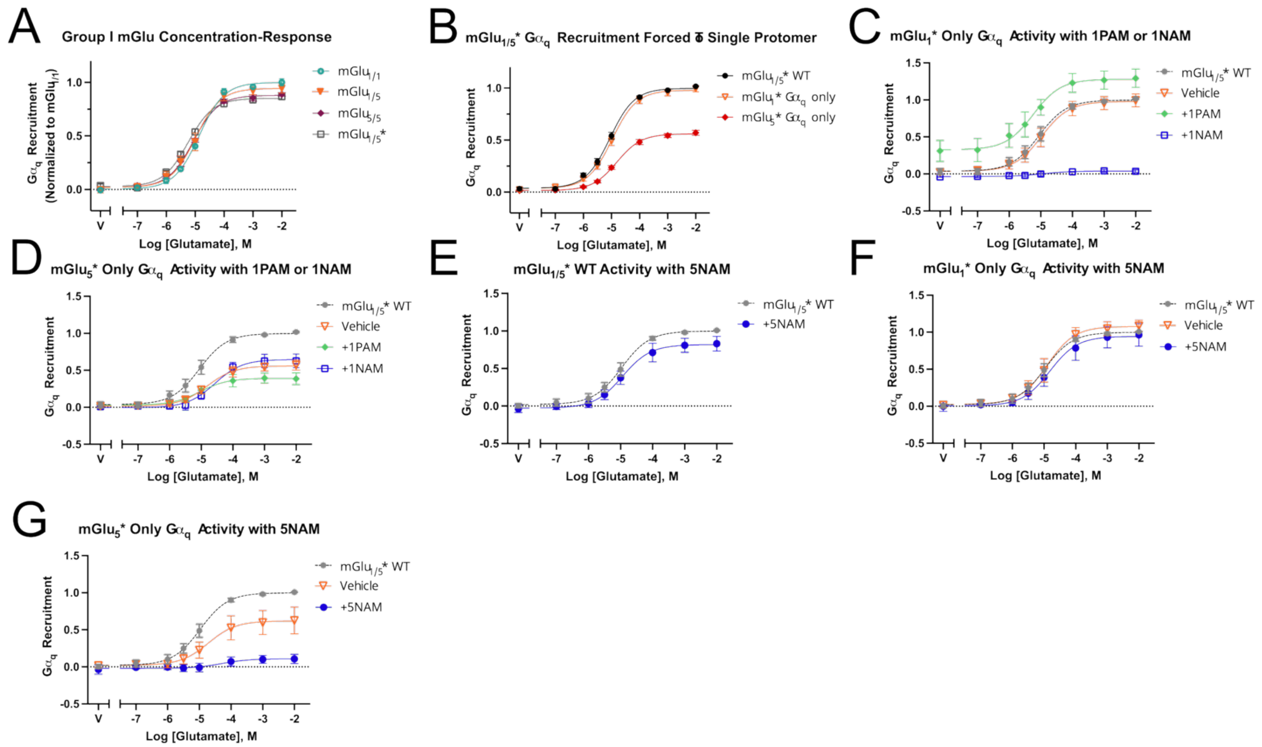

**Figure S5. Polarity Repeats of Key Experiments.**
mGlu$_1$-LgBiT and mGlu$_5$-HiBiT were used for all of the heterodimer experiments in the primary text due to very slight deficits in mGlu$_5$-LgBiT activity. Here are repeats of the key experiments using the heterodimer with mGlu$_1$-HiBiT/mGlu$_5$-LgBiT (mGlu$_{1/5}$*).
(A) Concentration-response curve from Figure 1 with mGlu$_{1/5}$* WT overlayed (black box).
(B) Concentration-response curve of mGlu$_{1/5}$* WT (black) versus mGlu$_1$/mGlu$_5^{F767D}$* (light orange) versus mGlu$_1^{F781D}$/mGlu$_5$* (dark orange) corresponding to Figure 4 in the main text.
(C) Concentration-response curve of mGlu$_{1/5}$* WT (grey) versus mGlu$_1$/mGlu$_5^{F767D}$* (orange) with PAM (100nM VU6024578-green) or NAM (100nM FITM-blue) corresponding to Figure 6.
(D) Concentration-response curve of mGlu$_{1/5}$* WT (grey) versus mGlu$_1^{F781D}$/mGlu$_5$* (orange) with PAM (100nM VU6024578-green) or NAM (100nM FITM-blue) corresponding to Figure 6.
(E) Concentration-response curve of mGlu$_{1/5}$* WT with vehicle (grey), with NAM (1 uM MTEP-blue).
(F) Concentration-response curve of mGlu$_{1/5}$* WT with vehicle (grey), mGlu$_1$/mGlu$_5^{F767D}$* with vehicle (orange), and mGlu$_1$/mGlu$_5^{F767D}$* with NAM (1uM MTEP-blue).
(G) Concentration-response curve of mGlu$_{1/5}$* WT with vehicle (grey), mGlu$_1^{F781D}$/mGlu$_5$* with vehicle (orange), and mGlu$_1^{F781D}$/mGlu$_5$* with NAM (1 uM MTEP-blue). Symbols represent the mean drug-induced BRET response and error bars represent ± SEM. The exact number of 'n' independent experiments and technical replicates are reported in Table S2.
